# Enhanced anti-nociception by novel dual antagonists for 5HT2AR and mGluR5 in preclinical models of pain

**DOI:** 10.1101/2025.05.21.655254

**Authors:** Daekyu Choi, Hyun Jin Heo, Haeyoung Shin, Jayzoon Im, Geonho Lee, Ah Hyun Kim, Kwang-Hyun Hur, Yoonmi Nho, Choon-Gon Jang, Hanmi Lee

## Abstract

Significant research has focused on developing anti-nociceptive pain treatments by targeting specific molecular candidates. The serotonin 2A receptor (5-HT2AR) and metabotropic glutamate receptor 5 (mGluR5) are recognized as key mediators in neuropathic pain. However, the combined effects of simultaneous inhibition of these targets remain unexplored. This current study investigated the therapeutic potential of concurrently antagonizing 5-HT2AR and mGluR5. Using spinal nerve ligation (SNL) and formalin-induced pain models in male Sprague-Dawley rats, we demonstrated that the simultaneous administration of both antagonists significantly enhanced anti-allodynic and anti-nociceptive effects, resulting in increased allodynia thresholds and reduced pain-related behaviors. This dual antagonism provided pain relief comparable to that of gabapentin and morphine. Furthermore, novel small molecules designed to simultaneously antagonize 5-HT2AR and mGluR5 exerted anti-nociceptive effects by suppressing excitatory postsynaptic responses and inhibiting the phosphorylation of ERK and AKT signaling molecules. Notably, the dual antagonist maintained anti-allodynic efficacy with repeated administration, unlike morphine, which exhibited clear tolerance development with daily use. Moreover, when administered intravenously, the dual antagonist demonstrated a low potential for abuse. These findings indicate that the simultaneous antagonism of 5-HT2AR and mGluR5 represents a promising pharmacological target for the management of chronic pain. This approach may offer enhanced analgesic outcomes while reducing the risk of undesirable side effects, such as tolerance and the potential for abuse.

## 1. Introduction

Extensive research has focused on identifying novel drug targets for the treatment of complex diseases, such as pain. Among the candidate targets, the serotonin (5-HT) and glutamate systems have garnered considerable interest due to their widespread expression patterns and involvement in various multifactorial diseases ^1,2^. Beyond their roles in the sleep/wake cycle and mood disorders (e.g., anxiety, depression, schizophrenia), 5-HT is increasingly recognized for its role in pain signaling. 5-HT levels rise rapidly during inflammation or injury, potentially acting as a pro-algesic mediator ^1^. In the spinal cord, elevated levels of 5-HT have been observed following high-intensity stimuli applied to the sciatic nerve of cats ^3^, similar to the pattern seen during noxious stimuli. The serotonin 2A receptor (5-HT2AR), one of the serotonin receptor subtypes expressed in dorsal root ganglia (DRG) and the spinal cord ^4–6^, appears to play a pivotal role in pain transmission. Increased expression of 5-HT2AR has been observed in the spinal cord dorsal horn following vincristine-induced neuropathy ^5^. Injection of 5-HT2AR agonists enhances pain responses induced by heat, mechanical stimuli, and inflammatory agents such as carrageenan, while pretreatment with 5-HT2AR antagonists like ketanserin attenuates these responses ^6–8^. The activation of 5-HT2AR by 5-HT is implicated in the regulation of neuronal excitability ^9,10^, consistent with increased c-Fos expression, a marker of neuronal activity following allodynic stimuli ^6^. These findings collectively highlight the significant role of 5-HT2AR in modulating pain signaling pathways. The metabotropic glutamate receptor 5 (mGluR5) is widely expressed throughout the brain, including regions such as the hippocampus, amygdala, cortex, midbrain, and spinal cord ^11^. Expression of mGluR5 increases following sciatic nerve injury, predominantly in the superficial layers of the spinal cord ^11,12^. Activation of mGluR5 with agonists, such as dihydroxyphenylglycine (DHPG), induces mechanical allodynia and hyperalgesia response to various stimuli including heat, cold, and mechanical touch ^13^. Conversely, mGluR5 antagonists elevate pain threshold ^14,15^. It has been reported that mGluR5 plays a crucial role in nociceptive glutamate signaling by regulating the release of glutamate from presynaptic terminals ^16,17^.

Although substantial evidence suggests that 5-HT2AR or mGluR5 are promising drug targets for neuropathic pain, no successful therapies have yet been translated into human applications. This lack of translation is primarily attributed to insufficient efficacy and/or intolerable adverse effects observed in human trials. Multi-target drug approaches have emerged as promising strategies to overcome the limitations of pain medications, partly owing to their potential for enhanced therapeutic effects and reduced toxicity. Previously, Pang et al. (2012) demonstrated a case study identifying a nociceptive multi-target drug, VVZ-149 (Opiranserin), which simultaneously inhibits 5-HT2AR and glycine transporter type 2 (GlyT2). Preclinical studies in various animal models of pain have shown that VVZ-149 exerts potent anti-allodynic and anti-nociceptive effects through its dual antagonism^18^. These promising results have been further supported by the recent Phase 3 clinical trial, which validated the efficacy of VVZ-149 in alleviating postoperative pain in human subjects ^19–22^.

In this study, we present compelling evidence demonstrating the enhanced effects of dual antagonism of both 5HT2AR and mGluR5 to achieve enhanced anti-allodynic and anti-nociceptive efficacy. Furthermore, we propose that a dual antagonist targeting both 5-HT2AR and mGluR5 holds strong potentials as a novel therapeutic agent for pain management.

## 2. Results

### 2.1 Enhanced effects of mGluR5 and 5-HT2AR antagonism alleviates the neuropathic pain

Previous studies have demonstrated the individual roles of 5-HT2AR and mGluR5 antagonists in producing anti-allodynic and anti-nociceptive effects ^9,12,17,18,23,24^. However, any interactions between these two receptors in alleviating neuropathic pain remains unexplored. We investigated the combined effects of 5-HT2AR and mGluR5 antagonism in two well-established rat models of pain: SNL and the formalin-induced hyperalgesia model ^25,26^ (**Fig. 1**).

**Figure 1.**
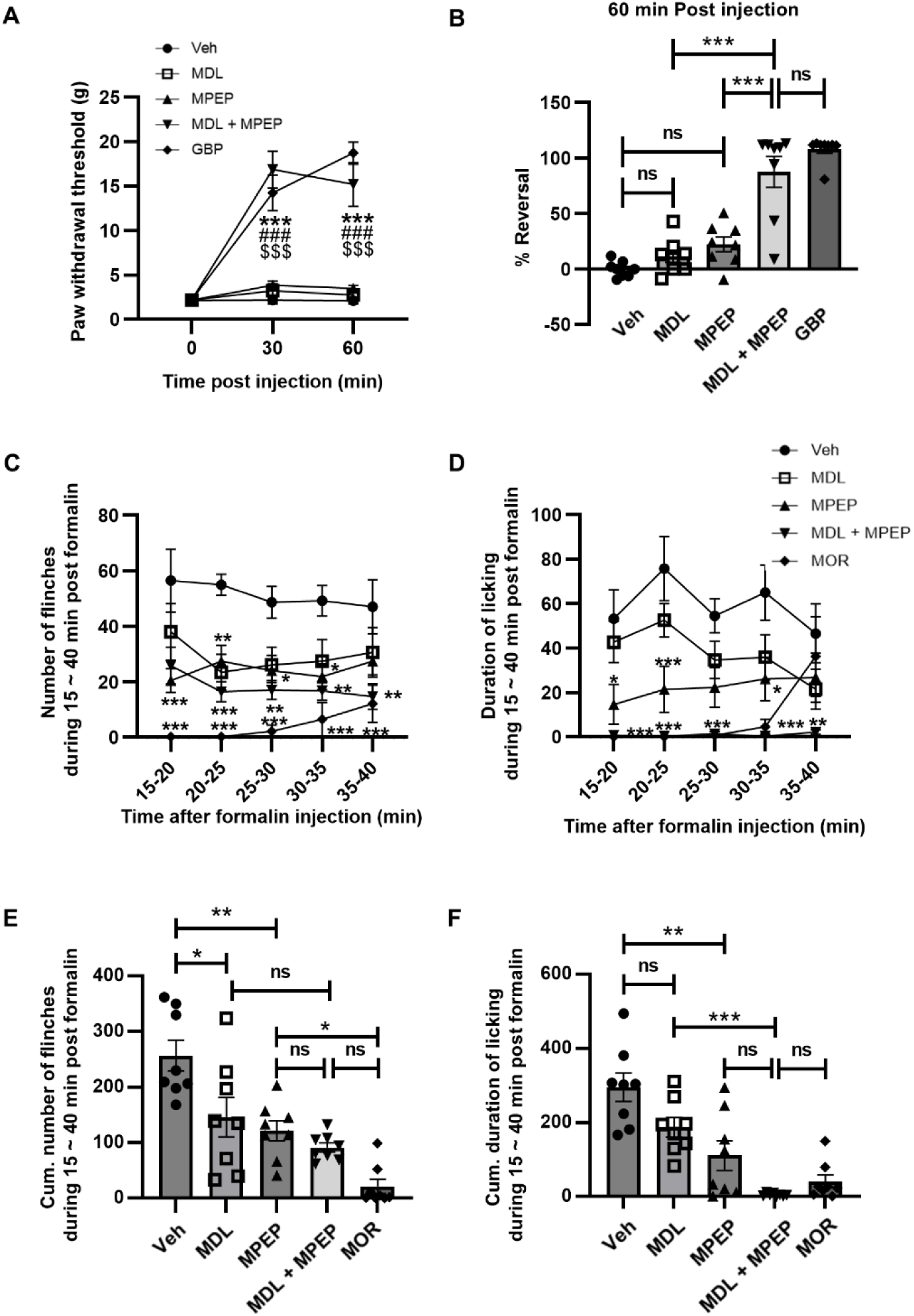
Enhanced effects of dual antagonism for mGluR5 and 5-HT2AR. **(A, B)** Anti-allodynic effects of individual or a combination of MDL11,939 and MPEP. **(A)** Withdrawal thresholds to mechanical stimulations of ipsilateral hind paw **(B)** Mean percent (%) reversal of the paw withdrawal thresholds 60-minute following treatment of Veh (vehicle) (2:8 DMA/PG, s.c.), MDL11,939 (MDL, 5 mg/kg, s.c.), MPEP (10 mg/kg, s.c.), a combination of MDL and MPEP, and gabapentin (GBP, 65 mg/kg, i.p.) in the SNL animal model. \*\*\**P* < 0.001 *vs* Vehicle; ###*P* < 0.001 *vs* MDL; $$$*P* < 0.001 *vs* MPEP, Two-way ANOVA. (N = 8 rats per group). **(C-F)** Anti-nociceptive effects of individual or a combination of MDL and MPEP in the formalin-induced pain model. **(C, D)** Time-course of flinches **(C)** and licking **(D)** responses induced by formalin injection to the rats treated with vehicle, MDL, MPEP, a combination of MDL and MPEP, or Morphine (MOR, 2 mg/kg, s.c.). \**P* < 0.05; \*\**P* < 0.01; \*\*\**P* < 0.001 *vs* Vehicle, Two-way ANOVA. **(E)** The cumulative number of flinches and **(F)** cumulative time duration of licking behavior for 15–40 minutes after the formalin injection. \**P* < 0.05; \*\**P* < 0.01; \*\*\**P* < 0.001, One-way ANOVA. (N = 8 rats per group). Error bars present S.E.M., “ns” means no significance (*P* > 0.05).

In the SNL model (**Fig. 1A, B**), the threshold for mechanical allodynia was investigated by measuring PWTs using a series of von Frey filaments. The PWTs were converted to % Reversal to illustrate the degree of recovery ^25^. Rats administered gabapentin (GBP, 65 mg/kg), a reference drug for the treatment of neuropathic pain, exhibited an 88% increase in % Reversal 60 minutes post-administration, indicating a substantial elevation in PWTs in response to mechanical stimuli. It was demonstrated that high doses of mGluR5 or extremely potent 5-HT2AR antagonists can reduce mechanical pain sensation ^12,27^. On the other hand, individual treatment with low doses of the selective 5-HT2AR antagonist, MDL11,939 (5 mg/kg) ^28,29^, or the mGluR5 antagonist, MPEP (10 mg/kg), resulted in only a modest increase in % Reversal (20 to 30%) compared to the vehicle group (n.s., *P* > 0.05) (**Fig.1B**). However, co-administration of MDL11,939 (5 mg/kg) and MPEP (10 mg/kg) significantly enhanced % Reversal (\*\*\**P* < 0.001), reaching levels comparable to those of gabapentin at 60-minute post-administration (*P* > 0.05 *vs* GBP). To further explore the potential involvement of other receptors, we investigated the effects of combining MDL11,939 (5 mg/kg) with radiprodil (30 mg/kg), a specific antagonist of NR2B-containing NMDA receptors, which are also implicated in neuropathic pain ^30,31^. However, this combination did not further enhance % Reversal compared to either drug alone and resulted in a significantly lower % Reversal than gabapentin (\*\*\**P* < 0.001 *vs* GBP) (**Fig. S1**).

Formalin-induced inflammation is known to cause both acute and tonic phase of pain, resulting in characteristic flinching and licking behaviors ^26^. Notably, the second phase, emerging approximately 10 minutes after formalin injection, is believed to be driven by central sensitization mechanisms within the dorsal horn of the spinal cord ^32–35^. Given this, the formalin model serves as a suitable paradigm for investigating central sensitization, and in our study, we specifically analyzed the effect of VVZ-2471 on the late phase of the pain response.

Consistent with the previous studies, subcutaneous formalin injection elicited significant flinching and licking behaviors, which were markedly suppressed by morphine (2 mg/kg) ^32,33^. Treatment with either MDL11,939 (10 mg/kg) or MPEP (5 mg/kg) individually reduced the number of flinches and the duration of licking time compared to the vehicle. However, their combined administration appeared to enhance these reductions **(Fig, 1C, D)**. To quantitatively assess the extent of this effect, the cumulative number of flinches and licking duration were analyzed during the 15–40 minutes period following drug administration. The results indicated a 40–60% reduction in cumulative number of flinches by MDL11,939 or MPEP compared to the vehicle control (\**P* < 0.05, \*\**P* < 0.01). Despite this, the combination treatment did not further decrease the number of flinches beyond what was observed with MDL11,939 or MPEP alone (n.s., *P* > 0.05) (**Fig. 1E**). In contrast, the combined antagonist treatment significantly reduced licking duration compared to MDL11,939 treatment alone (\*\*\**P* < 0.001), exhibiting an effect comparable to that of morphine (n.s., *P* > 0.05) (**Fig. 1F**). This suggests that the simultaneous antagonism of 5-HT2AR and mGluR5 can effectively exert anti-allodynic and anti-nociceptive effects, alleviating nociceptive pain. Overall, the significant analgesic effects observed with low doses of MDL11,939 and MPEP in both the SNL and formalin-induced pain models indicate that the combination of 5-HT2AR and mGluR5 antagonists may improve therapeutic effects in pain treatment. These findings further underscore mGluR5 as a promising target for combination therapy alongside 5-HT2AR antagonists.

### 2.2 Development of dual antagonists for 5-HT2AR and mGluR5

We synthesized over 1,000 chemical compounds with potential dual antagonistic activity against mGluR5 and 5-HT2AR, demonstrating high selectivity and moderate potency. Among these, VVZ-2471 and VVZ-2868 were identified as promising candidates for dual antagonist therapy. VVZ-2471 demonstrated IC_50_ values of 438.1 nM for human 5-HT2AR and 87.1 nM for human mGluR5 (**Fig. 2A and B**). Similarly, VVZ-2868 exhibited IC_50_ values of 615 nM for human 5-HT2AR and 156 nM for human mGluR5 (data not shown). Notably, VVZ-2471 exhibited a two-fold higher potency for mGluR5 compared to VVZ-2868. VVZ-2471 exhibited higher IC_50_ values for 5-HT2BR and 5-HT2CR (1.7 and 3.8 μM, respectively), indicating lower potency for these receptors compared to 5-HT2AR. Similarly, functional studies for mGluR1, mGluR7, and mGluR8 demonstrated IC_50_ values exceeding 10 μM (**Fig. 2B**). These data suggest that VVZ-2471 preferentially targets 5-HT2AR and mGluR5.

**Figure 2.**
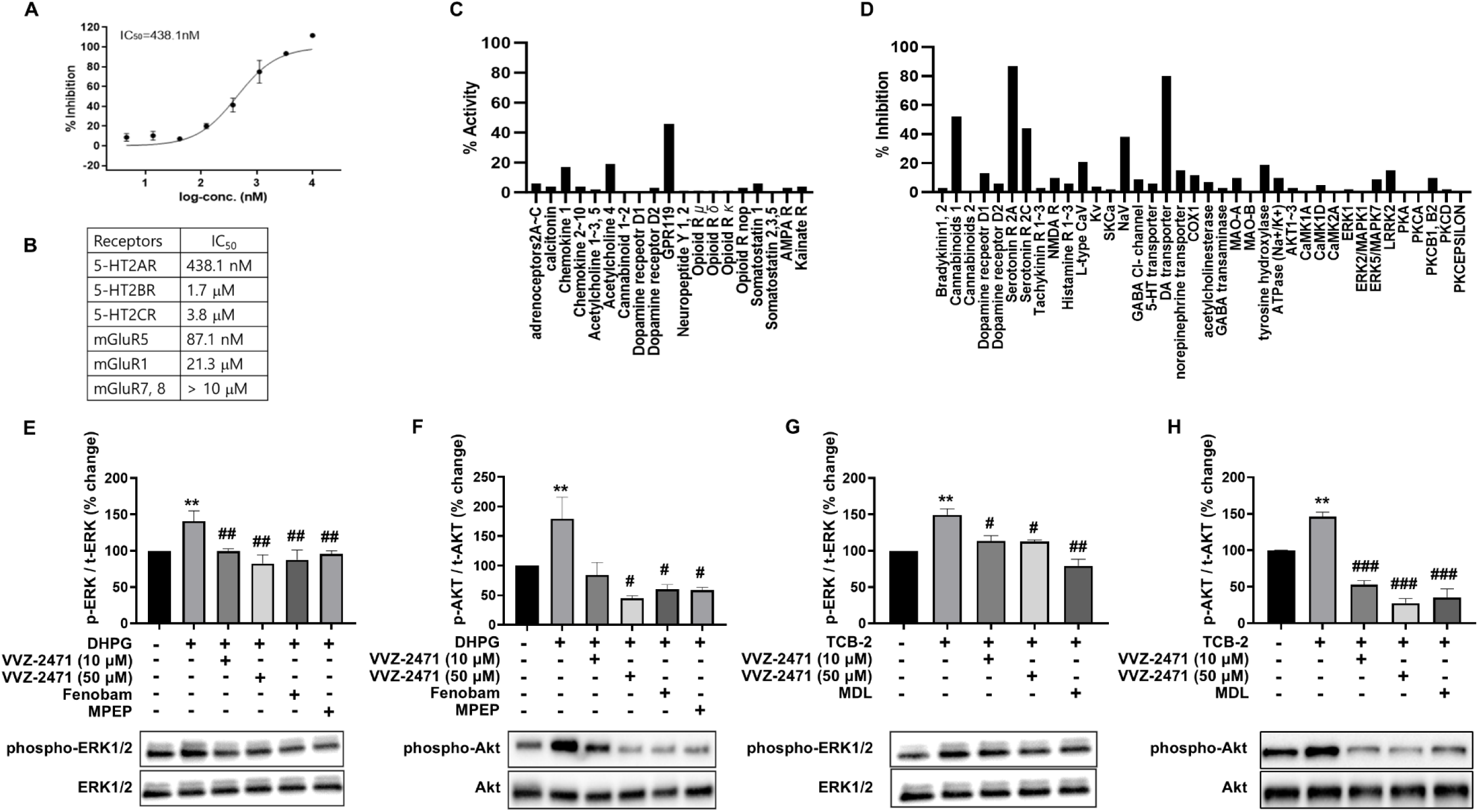
Dual antagonistic activities of VVZ-2471 for 5-HT2AR and mGluR5. **(A)** IC_50_ for the accumulation of IP1 *via* 5-HT2AR activation. **(B)** The selectivity profile of VVZ-2471. **(C, D)** Target specificity of VVZ-2471 at 10 μM among 168 GPCRs, 371 Kinases, 87 ion channels, transporters, and enzymes. **(C)** % activities and **(D)** % inhibition of VVZ-2471 on molecules related to the pain pathway. No significant activity (≥ 50%) except for dopamine reuptake transporter (DAT, 80% inhibition) and cannabinoid receptor 1 (CB1, 52% inhibition). **(E–H)** Inhibition of p-ERK1/2 and p-AKT increments by VVZ-2471 in cerebral cortical neurons in Western blot. **(E–F)** Increase of p-ERK1/2 **(E)** and p-AKT **(F)** induced by DHPG (1 μM) were inhibited by VVZ-2471 (10 and 50 μM). Fenobam (50 μM) and MPEP (50 μM) as references of mGluR5 antagonists (N = 3 cultures per group). **(G–H)** Changes of p-ERK1/2 **(G)** and p-AKT **(H)** induced by TCB-2 (0.1 and 10 μM, respectively) in the presence of either VVZ-2471 (10 and 50 μM) or MDL (MDL11,939) (50 μM) (N = 3 cultures per group). Error bars present S.E.M., % changes normalized to the no treatment condition., \*\**P* < 0.01 *vs* non-treated; #*P* < 0.05, ##*P* < 0.01 *vs* either DHPG or TCB-2 only, Two-tailed t-test.

To further investigate the selectivity of VVZ-2471, multiple screening experiments were conducted encompassing GPCRs, enzymes, ion channels, and various kinases, using 10 μM of VVZ-2471. This comprehensive screening aimed to identify any potential interactions of VVZ-2471 with targets other than 5-HT2AR and mGluR5. Among the potential targets involved in pain pathways ^36,37^, VVZ-2471 exhibited a weak agonist effect on GPR119, with an effect less than 50% of that of the reference compound. Additionally, it displayed weak inhibitory potential against the dopamine reuptake transporter (DAT), as well as 5-HT2CR. The impact on other targets remained minimal (**Fig. 2C and D**). These results indicate a high specificity of VVZ-2471 for 5-HT2AR and mGluR5, with minimal effects on other major targets implicated in pain pathways such as opioid receptors (*mu-, delta-, kappa-, and opioid-related nociceptive receptor*), NMDA receptor, GABAA receptor, cannabinoid receptors (1 and 2), and voltage gated Ca^2+^ channels ^37^.

### 2.3 Pharmacokinetic profile of VVZ-2471 supports its suitability for oral pain treatment

To evaluate the suitability of VVZ-2471 as an orally administered drug, we investigated its systemic exposure in male rats. VVZ-2471 was administered *via* oral routes (**Table 1**). After a single oral administration of VVZ-2471 (10 mg/kg) to male rats, the key pharmacokinetic parameters were as follows: *t_1/2_* of 1.53 hours, *T_max_* ranging from 0.5 to 2 hours, and *C_max_* of 1731 ng/mL. These values indicate that VVZ-2471 is absorbed quickly and effectively after oral administration. VVZ-2471 demonstrated high brain penetration. Following a single oral administration of 50 mg/kg, the brain-to-plasma ratio based on AUC_0-last_ was 1.87 (measured at 2 hours post-administration). This ratio indicates that VVZ-2471 readily crosses the blood-brain barrier and achieves significant concentrations in the brain, classifying it as a drug with high brain tissue distribution ^38^. Oral administration of VVZ-2471 at various doses (10, 25, and 50 mg/kg) resulted in a dose-dependent increase in plasma concentration. The plasma protein binding of VVZ-2471 was high in humans (98.7%), rats (98.1%), and dogs (96.6%). The assessment of metabolic stability revealed a shorter *t*_1/2_ in rat liver microsomes (80 minutes) compared to dogs and humans (212.2 minutes and 417.6 minutes, respectively). These findings collectively suggest that VVZ-2471 possesses the pharmacokinetic properties favorable for an oral pain treatment, including high systemic exposure, efficient blood-brain barrier penetration, and substantial plasma protein binding.

**Table 1.**
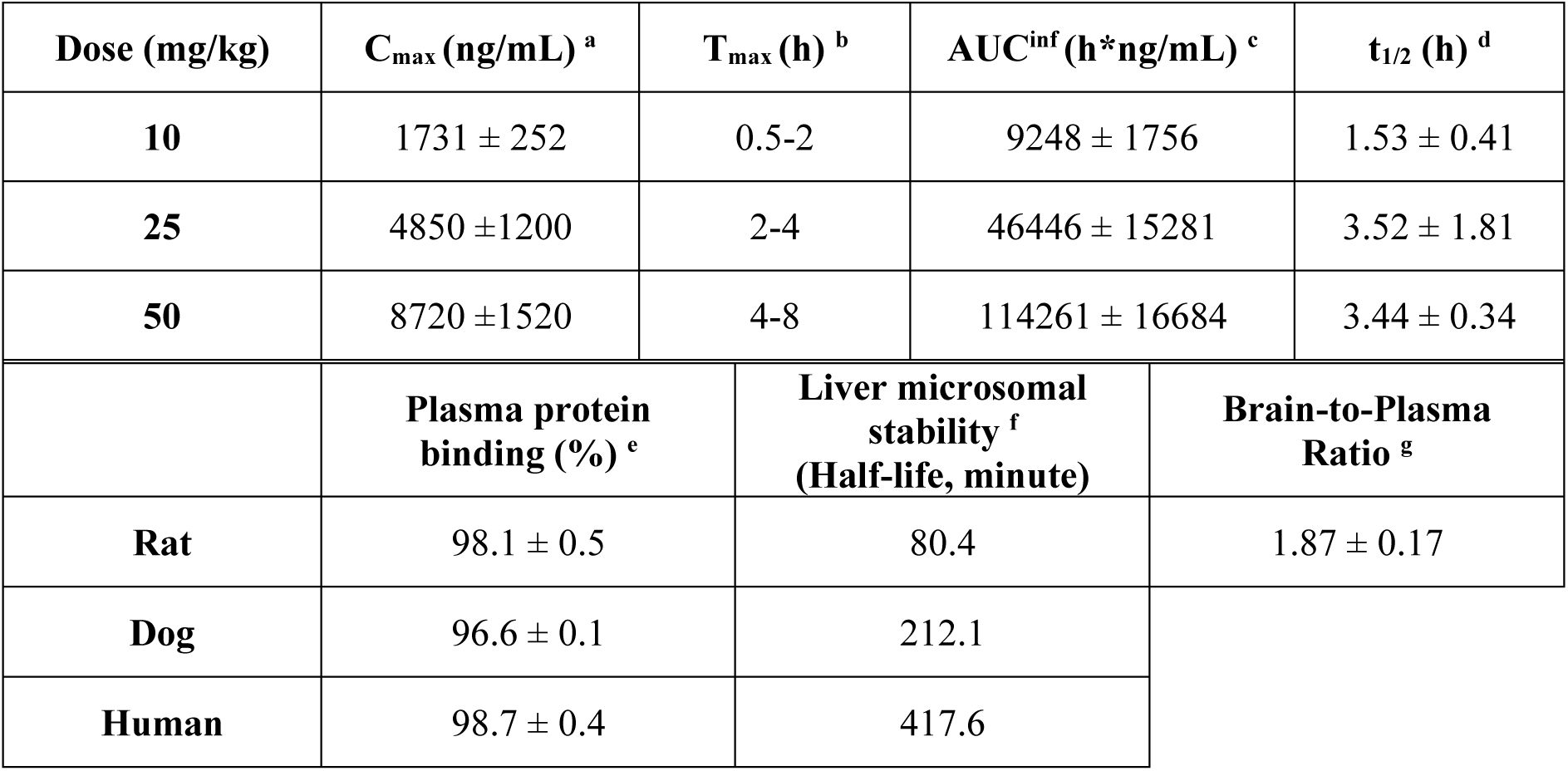
Pharmacokinetic parameters were obtained using noncompartmental analysis by Pheonix 64 WinNonlin. Data presented as mean ± S.D. T_max_ suggested as range. **^a^** Maximum observed concentration (C_max_), **^b^** Time to maximum observed concentration (T_max_), **^c^** Area under the concentration-time curve from time zero to the extrapolated area from the last measurable concentration to infinity (AUC_inf_), **^d^** Terminal elimination half-life (t_1/2_), **^e^** % protein bound fraction of VVZ-2471 (10 µM) in plasma for 4 hours reaction time, **^f^** Half-life of VVZ-2471 (1 µM) in the presence of 0.5 mg/mL of protein concentration of liver microsomes during 1 h incubation, **^g^** Ratio of VVZ-2471 exposure in brain vs plasma calculated from AUC from 0 to 2 h post-dosing (50 mg/kg). (n=3 rats per dose or test).

| Dose (mg/kg) | C<sub>max</sub> (ng/mL) <sup>a</sup> | T<sub>max</sub> (h) <sup>b</sup> | AUC<sup>inf</sup> (h*ng/mL) <sup>c</sup> | t<sub>1/2</sub> (h) <sup>d</sup> |
| --- | --- | --- | --- | --- |
| **10** | 1731 ± 252 | 0.5-2 | 9248 ± 1756 | 1.53 ± 0.41 |
| **25** | 4850 ± 1200 | 2-4 | 46446 ± 15281 | 3.52 ± 1.81 |
| **50** | 8720 ± 1520 | 4-8 | 114261 ± 16684 | 3.44 ± 0.34 |
|  | **Plasma protein binding (%) <sup>e</sup>** | **Liver microsomal stability <sup>f</sup>**<br>**(Half-life, minute)** | **Brain-to-Plasma Ratio <sup>g</sup>** |  |
| **Rat** | 98.1 ± 0.5 | 80.4 | 1.87 ± 0.17 |  |
| **Dog** | 96.6 ± 0.1 | 212.1 |  |  |
| **Human** | 98.7 ± 0.4 | 417.6 |  |  |

### 2.4 The dual antagonist inhibits synaptic functions mediated by 5-HT2AR and mGluR5

It is well known that the phosphorylation of ERK1/2 and AKT, which are downstream molecules of mGluR5 and 5-HT2AR, plays a crucial role in regulating neuropathic pain and its perception ^39–41^. To investigate the functional effects of VVZ-2471 on neurons, an *in vitro* study was conducted using cerebral cortical neurons. These neurons were stimulated with the mGluR5 agonist DHPG, both in the absence and presence of VVZ-2471 (**Fig. 2E ∼ H**). The resulting phosphorylation levels (% changes) of ERK1/2 and AKT induced by DHPG in the presence of VVZ-2471 (10 and 50 μM), fenobam, or MPEP were compared to those observed with DHPG alone. In addition, % changes in ERK1/2 and AKT phosphorylation levels were examined in groups treated with DHPG in the presence of Fenobam or MPEP ^39,42^. As expected, DHPG significantly increased ERK1/2 and AKT phosphorylation (\*\**P* < 0.05 *vs* untreated control). However, VVZ-2471 effectively inhibited DHPG-induced phosphorylation, similar to the effects observed with MPEP and Fenobam (#*P* < 0.05, ##*P* < 0.01 *vs* DHPG alone) (**Fig. 2E and F**). Similarly, the 5-HT2AR specific agonist TCB-2 also increased ERK1/2 and AKT phosphorylation (\*\**P* < 0.05 *vs* untreated control), which was significantly reduced by VVZ-2471 (#*P* < 0.05, ###*P* < 0.001 *vs* TCB-2 alone) (**Fig. 2G and H**). These findings demonstrate that VVZ-2471 effectively inhibits both mGluR5 and 5-HT2AR-mediated signaling responses in neurons by suppressing phosphorylation of ERK1/2 and AKT.

Several studies indicate that enhanced activity of 5-HT *via* 5-HT2AR ^9,43,44^ and glutamate *via* mGluR5 contributes to lower thresholds to detect pain by increasing excitability ^16,17,45^ and reducing pain thresholds ^12^, which result in hypersensitivity to allodynic stimuli. Therefore, we examined the potential impact of the dual antagonist on synaptic activity using *ex vivo* slice preparations of rat spinal cord and investigated whether VVZ-2471 functionally inhibits mGluR5 and 5-HT2AR.

Spontaneously generated EPSCs (sEPSC) were measured in neurons located within the superficial dorsal horn, which was visually identified as a translucent region (**Fig. 3A**). The application of DHPG (3 μM) significantly increased the frequency of sEPSCs (\*\*\**P* < 0.001 *vs* control). However, VVZ-2471 (10 μM) completely inhibited the DHPG-induced increase (**Fig. 3B ∼ D**), similar to the effects observed with MPEP (10 μM) and Basimglurant (10 μM) (*P* > 0.05, **Fig. S2**). These results suggest that VVZ-2471 effectively prevents mGluR5-mediated enhancement of synaptic activity in the spinal cord, similar to the effects of MPEP and Basimglurant, which are well-known selective and potent mGluR5 antagonists.

**Figure 3.**
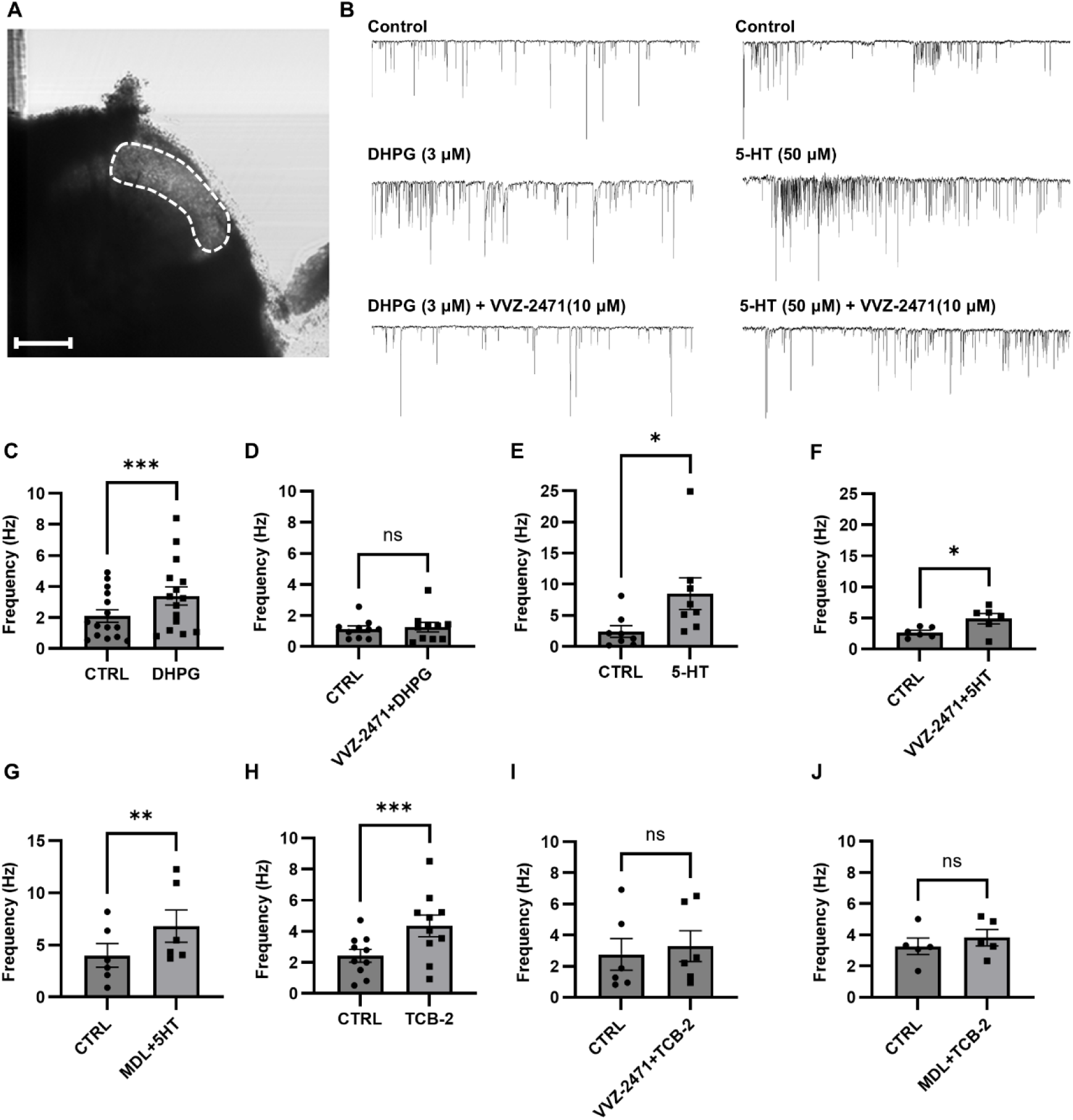
Inhibition of synaptic responses by a dual antagonist, VVZ-2471. **(A)** The Image of the superficial dorsal horn of the spinal cord (Scale bar, 200 μm). **(B)** Example traces for sEPSC induced by either 5-HT (Right, 50 μM) or DHPG (Left, 3 μM) in the absence and presence of VVZ-2471 (10 μM). **(C)** Frequency (Hz) change of sEPSC induced by DHPG (Control (CTRL): 2.1 ± 0.38; DHPG: 3.4 ± 0.57; n=15 / N=9) **(D)** Frequency (Hz) change of sEPSC induced by DHPG in the presence of VVZ-2471 (CTRL: 1.12 ± 0.20; DHPG + VVZ-2471: 1.24 ± 0.33, n=9 / N=3). **(E)** Frequency (Hz) changes induced by 5-HT (CTRL: 2.4 ± 0.88; 5-HT: 8.50 ± 2.38, n=8 / N=6). **(F)** Frequency (Hz) changes induced by 5-HT in the presence of VVZ-2471 (CTRL: 2.68 ± 0.30; 5-HT + VVZ-2471: 4.89 ± 0.77, n=6 / N=3). **(G)** Frequency (Hz) changes induced 5-HT in the presence of MDL11,939 (10 μM, CTRL: 4.0 ± 1.03; 5-HT + MDL11,939: 6.8 ± 1.42, n=6 / N=4). **(H)** Frequency (Hz) changes induced by TCB-2 (100 μM, CTRL: 2.4 ± 0.39; TCB-2: 4.35 ± 0.66, n=10 / N=5). **(I)** Frequency (Hz) changes induced by TCB-2 in the presence of VVZ-2471 (CTRL: 2.75 ± 0.93; TCB-2 + VVZ-2471: 3.28 ± 0.91, n=6 / N=3). **(J)** Frequency (Hz) changes induced by TCB-2 in the presence of MDL11,939 (CTRL: 3.26 ± 0.47; TCB-2 + MDL 11,939: 3.82 ± 0.47, n=5 / N=3). Error bars present S.E.M., “ns” means no significance (*P* > 0.05); \**P* < 0.05; \*\**P* < 0.01; \*\*\**P* < 0.001, Two-tailed t-test.

The application of 5-HT (50 μM) significantly increased the frequency of sEPSC (\**P* < 0.05 *vs* control, **Fig. 3B and E**). However, neither VVZ-2471 (10 μM) nor MDL11,939 (10 μM) completely inhibited the frequency increase induced by 5-HT (\**P* < 0.05, \*\**P* < 0.01 *vs* control, **Fig. 3F and G**), possibly due to 5-HT activating other 5-HT receptors beyond 5-HT2AR. To investigate this possibility, TCB-2 (100 μM), a selective 5-HT2AR agonist, was applied instead of 5-HT ^46^. Consequently, TCB-2 significantly increased the frequency of sEPSC (\*\*\**P* < 0.001 *vs* control, **Fig. 3H**), an effect that was completely inhibited by both MDL11,939 and VVZ-2471 (*P* > 0.05, **Fig. 3I and J**), confirming the specific antagonism of 5-HT2AR-mediated synaptic activity by VVZ-2471. Importantly, VVZ-2471 alone did not alter the frequency of sEPSC (**Fig. 3D, F, and I**). Furthermore, no significant changes in sEPSC amplitudes were observed under any experimental conditions (**Fig. S3**). Collectively, these findings demonstrate that VVZ-2471 effectively inhibits both 5-HT2AR and mGluR5-mediated synaptic activity and signaling responses in neurons.

### 2.5 Dual antagonist effects in spinal nerve ligation and formalin-induced pain models

Previous data indicated that the simultaneous antagonism of 5-HT2AR and mGluR5 may effectively increase the threshold for detecting allodynia and nociceptive stimuli (**Fig. 1**). Based on these findings, the anti-allodynic and anti-nociceptive effects of VVZ-2471 were evaluated in the SNL and formalin-induced pain models as in the previous experiments, following oral administration of the dual antagonists (**Table 1**). In the SNL model, VVZ-2471 significantly increased the paw withdrawal threshold and % Reversal at 60- and 120-minutes post-administration (\*\*\**P* < 0.001) (**Fig. 4A, B**). The analgesic efficacy of VVZ-2471 showed a dose-dependent increase, with an ED50 of 9.29 mg/kg at 120 min (**Fig. 4C**). To further validate the pharmacological effects of VVZ-2471 in relation to allodynic stimuli detection, the blood samples were analyzed for VVZ-2471 immediately after measuring pain intensities, establishing a PK/PD relationship. The concentration of VVZ-2471 increased with increasing % Reversal, resulting in EC_50_ of 1830 ng/mL. At 20 mg/kg, full efficacy was achieved, reaching steady-state plasma concentrations of 4048 ± 405 ng/mL (12.4 ± 1.2 μM) (**Fig. 4D, E**), demonstrating a strong correlation between plasma concentration of VVZ-2471 and % Reversal (**Fig. 4E**). VVZ-2868, an analog of VVZ-2471 with 50% reduced potency for mGluR5, also demonstrated a dose-dependent increase in the paw withdrawal threshold, exhibiting an ED_50_ of 7.67 mg/kg with a well-fit PK/PD correlation, similar to VVZ-2471 (**Fig. 4F ∼ H**). The efficacy of both VVZ-2471 and VVZ-2868 at 20 mg/kg was comparable to that of gabapentin at 65 mg/kg and lasted for 120 minutes following administration (**Fig. 4A, B and F, G**).

**Figure 4.**
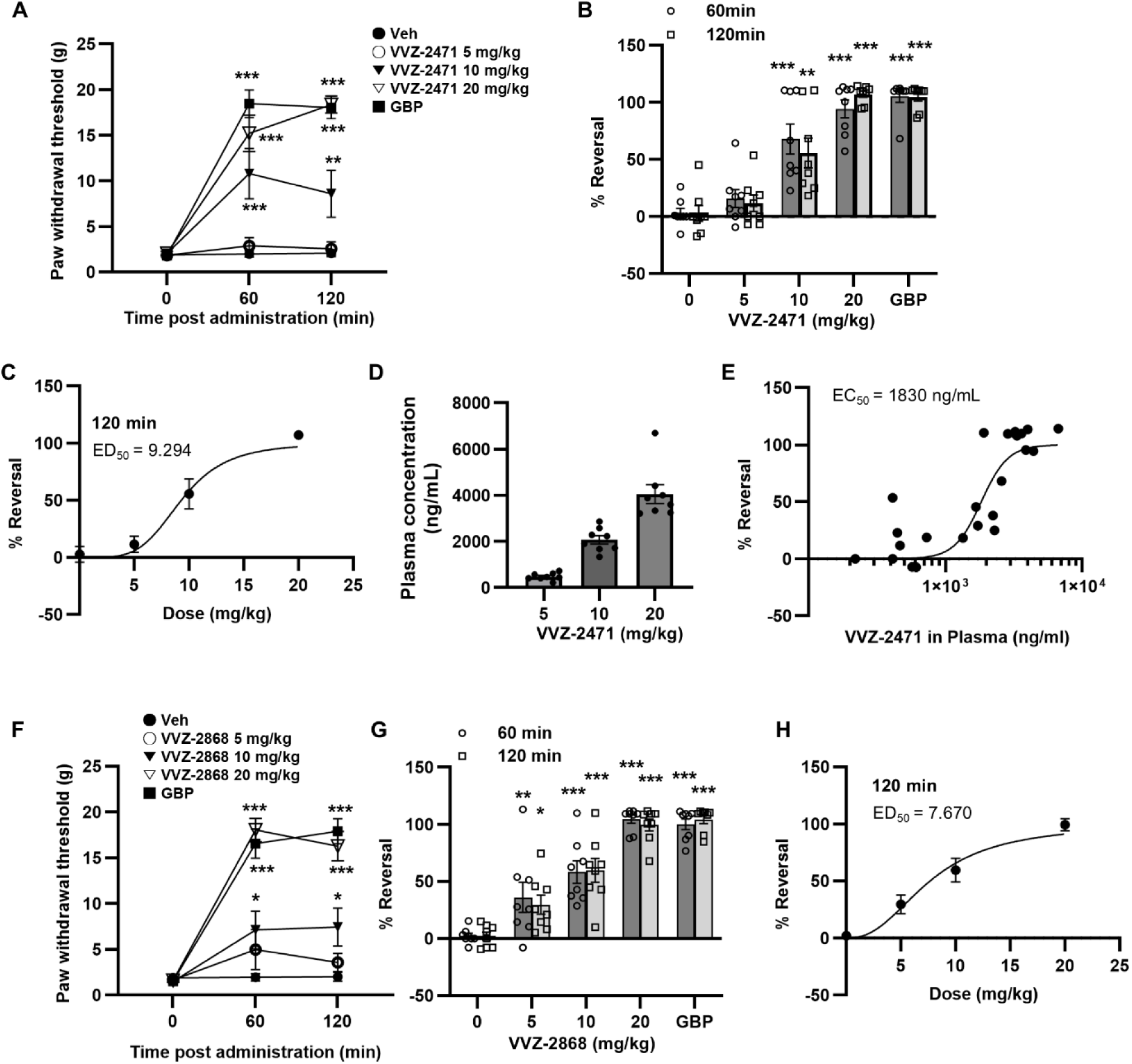
Anti-allodynic efficacy of the dual antagonists against 5-HT2AR and mGluR5 in the SNL model. **(A)** Withdrawal thresholds to mechanical stimulations of ipsilateral hind paw post administration of VVZ-2471. **(B)** Increased % reversal with escalating doses of VVZ-2471 (5, 10, 20 mg/kg, p.o.). **(C)** Dose-dependent % reversal following VVZ-2471 administration. The ED_50_ = 9.29 mg/kg at 120 minutes. **(D-E)** The PK/PD correlation of VVZ-2471. **(D)** Plasma concentrations (ng/mL) of VVZ-2471 at each dose (5 mg/kg, 484 ± 55; 10 mg/kg, 2077 ± 179; 20 mg/kg, 4048 ± 405). **(E)** % Reversals plotting to the corresponding plasma concentration of VVZ-2471. EC_50_ = 1830 ng/mL (5.66 µM). **(F)** Withdrawal thresholds to mechanical stimulations of ipsilateral hind paw post administration of VVZ-2868. **(G)** Increased % reversal with escalating doses of VVZ-2868 (5, 10, 20 mg/kg, p.o.). GBP as reference (65 mg/kg, i.p.). **(H)** Dose dependent % reversal following VVZ-2868 administration. ED_50_ = 7.67 mg/kg at **(I)** 120 minutes. N = 8 rats per group. Error bars present S.E.M., \*\**P* < 0.01; \*\*\**P* < 0.001 *vs* 0 mg/kg, Two-way ANOVA.

In the formalin-induced pain model, oral administration of VVZ-2471 and VVZ-2868 (100 minutes pretreatment) significantly reduced flinching numbers and licking time duration (\*\*\**P* < 0.001, \*\**P* < 0.01, \**P <* 0.05 *vs* Vehicle) (**Fig. S4**). At doses of 25 and 50 mg/kg, both dual antagonists showed a significant reduction in cumulative number of flinches and licking duration (\*\*\**P* < 0.001, \*\**P <* 0.01, \**P <* 0.05 *vs* Vehicle), achieving an efficacy comparable to morphine at 50 mg/kg (n.s., *P* > 0.05) (**Fig. 5A, B**). Further PK/PD correlation analysis revealed that VVZ-2471’s efficacy (% Control) corresponded to its blood concentration, yielding an EC_50_ of 4280 ng/mL (**Fig. 5C**). Our results strongly suggest that simultaneous antagonism against 5-HT2AR and mGluR5 may effectively exert analgesic efficacy in both the SNL and formalin-induced pain models.

**Figure 5.**
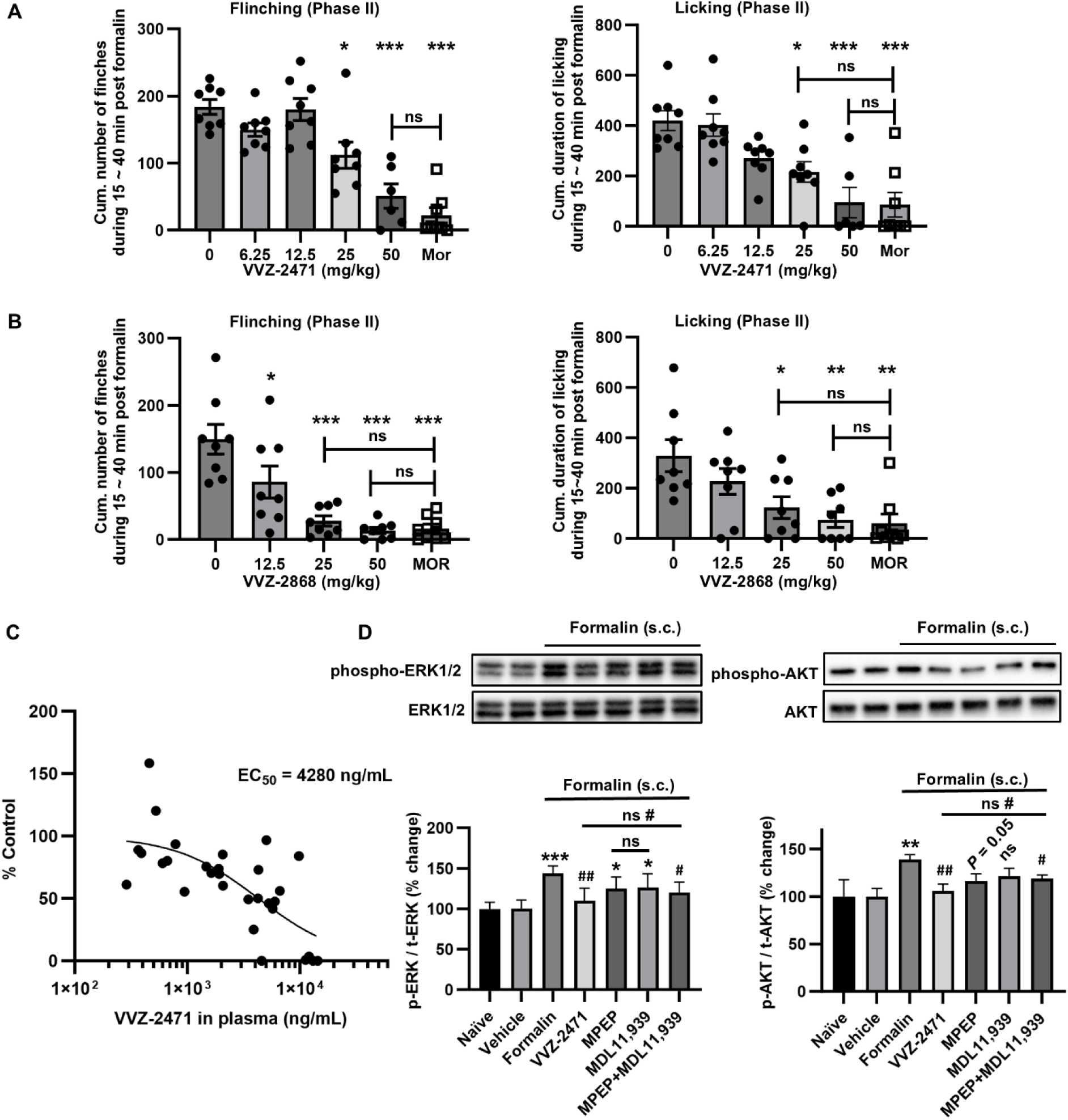
Anti-nociceptive efficacy of the dual antagonists against 5-HT2AR and mGluR5 mediated *via* ERK1/2 and AKT phosphorylation in the formalin-pain model. (**A, B**) Formalin-induced pain responses ameliorated by dual antagonists in rats. Cumulative flinch numbers and cumulative time duration of licking behavior for 15 to 40 minutes following administration of **(A)** VVZ-2471 (N = 8 each group except for N = 6 for 50 mg/kg, p.o.) and **(B)** VVZ-2868 (N = 8 rats per group, p.o.). Morphine (MOR, 2 mg/k, s.c.) as a reference. **(C)** The PK/PD correlation of VVZ-2471. % Controls plotting to the corresponding plasma concentration of VVZ-2471. EC_50_ was 4280 ng/mL (13.38 µM). **(D)** Increase of p-ERK1/2 and p-AKT by formalin was reduced by the dual antagonist of 5-HT2AR and mGluR5 (VVZ-2471, 50 mg/kg, p.o.). % changes normalized to naïve condition. MPEP (10 mg/kg, s.c.) and MDL11,939 (5 mg/kg, s.c.) (N=4 per group). Error bars present S.E.M., “ns” means no significance (P >0.05); \**P* < 0.05; \*\**P* < 0.01, \*\*\**P* < 0.001 *vs* vehicle group; #*P* < 0.05, ##*P* < 0.01 *vs* formalin group, One-way ANOVA for **(A)** and **(B)**; Two-tailed t-test for **(D)**.

In primary neuronal cultures, VVZ-2471 effectively suppressed the phosphorylation of ERK1/2 and AKT (**Fig. 2E ∼ H**). Therefore, we investigated whether the phosphorylation levels of ERK1/2 and AKT would be altered by VVZ-2471 *in vivo*. Following formalin-induced behavioral tests, spinal cords were isolated, and the levels of phosphorylation of ERK1/2 (p-ERK1/2) and AKT (p-AKT) were examined. Consistent with previous findings, formalin injection significantly increased p-ERK1/2 and p-AKT levels (\*\*\**P* < 0.001, \*\**P* < 0.01, Fig. 5D). However, VVZ-2471 (50 mg/kg) significantly reduced formalin-induced phosphorylation of both ERK1/2 and AKT (##*P* < 0.01 *vs* formalin). This reduction was also observed when both MPEP and MDL11,939 were administered simultaneously (#*P* < 0.05 *vs* formalin, *P* > 0.05 for VVZ-2471 *vs* MPEP + MDL11,939) (**Fig. 5D**). Single treatments with either MPEP or MDL11,939 did not result in significant reduction in the levels of p-ERK1/2 and p-AKT (*P* ≥ 0.05 *vs* formalin).

Collectively, our results provide compelling evidence that dual antagonists targeting mGluR5 and 5-HT2AR can effectively reduce allodynia and pain intensity, potentially through the modulation of downstream signaling molecules such as p-ERK1/2 and p-AKT.

### 2.6 The dual antagonist demonstrates no tolerance and low abuse liability in rats

It is well-known that the prolonged administration of analgesic drugs, particularly opioids, can lead to reduced efficacy due to the development of drug tolerance ^47^. To evaluate the potential for tolerance associated with VVZ-2471, we investigated its analgesic efficacy following repeated administrations in the SNL model. Over a 14-day daily administration period, VVZ-2471 (25 mg/kg) consistently maintained anti-allodynic effects, showing no significant difference between Day 1 and Day 14 (*P* > 0.05, **Fig. 6**). In contrast, morphine (3 mg/kg) exhibited a gradual reduction in efficacy over the same period, with over 80% loss of the initial effect by Day 14 (##*P* < 0.01, **Fig. 6**). These results confirm that VVZ-2471 does not induce tolerance in this rodent model of neuropathic pain.

**Figure 6.**
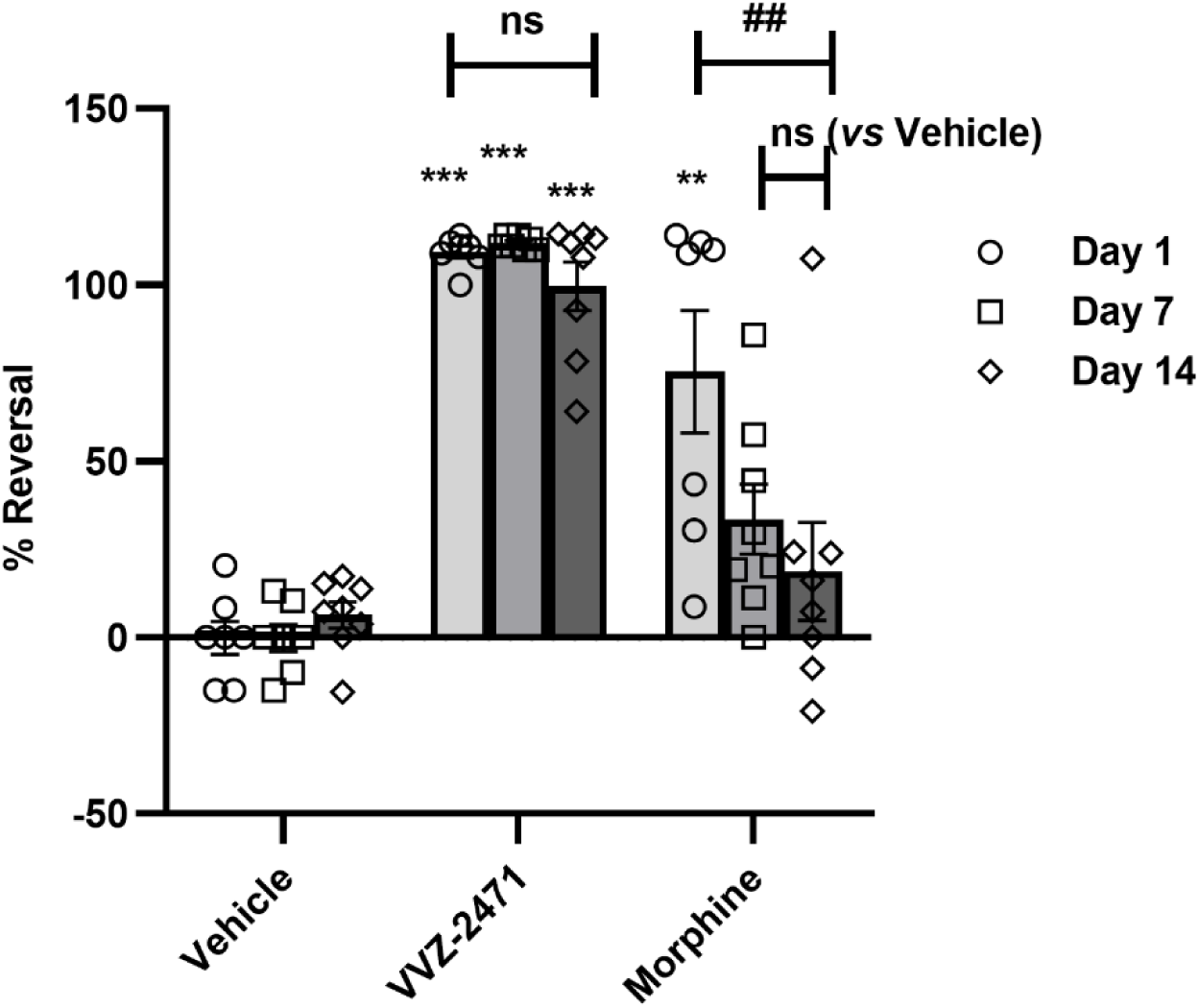
No tolerance of anti-allodynic effect of VVZ-2471. % Reversal measured on Day 1, Day 7, and Day 14. VVZ-2471 (25 mg/kg, p.o.) and morphine (3 mg/kg, s.c.) were administered daily (N=8 rats per group). Error bars present S.E.M., “ns” means no significance (*P* > 0.05); \*\**P* < 0.01, \*\*\**P* < 0.001 *vs* vehicle; ##*P* < 0.05 between Day 1 and Day 14 within the morphine group, Two-way ANOVA.

The reinforcing properties of a substance are intrinsically linked to its abuse liability and play a pivotal role in promoting drug-seeking behavior ^48^. To assess the abuse potential of VVZ-2471, we employed an intravenous self-administration (IVSA) model in rats (**Fig. 7A**), where the animals were trained to self-administer drugs by pressing levers in an operant conditioning chamber, with active lever presses serving as a direct measure of drug reinforcement. Morphine, a positive control, significantly increased both the number of infusion and active lever responses (\*\*\**P* < 0.001, **Fig. 7B**), consistent with its well-documented reinforcing properties. In contrast, VVZ-2471 did not produce a significant increase in self-administration, even at the highest dose (1.0 mg/kg/infusion), indicating a lack of reinforcing efficacy. Furthermore, morphine administration also elevated inactive lever presses, potentially indicative of drug-induced impulsivity, an effect absent following VVZ-2471 administration (**Fig. 7B**, bottom). Collectively, these findings suggest that VVZ-2471 possesses minimal reinforcing properties and, by extension, a low risk of inducing drug dependency. In summary, our findings provide compelling evidence that the simultaneous antagonism of 5-HT2AR and mGluR5 offers promising non-opioid analgesic solutions for chronic neuropathic pain. This approach demonstrates several advantages over morphine and gabapentin, including low potential for abuse and a reduced risk of tolerance.

**Figure 7.**
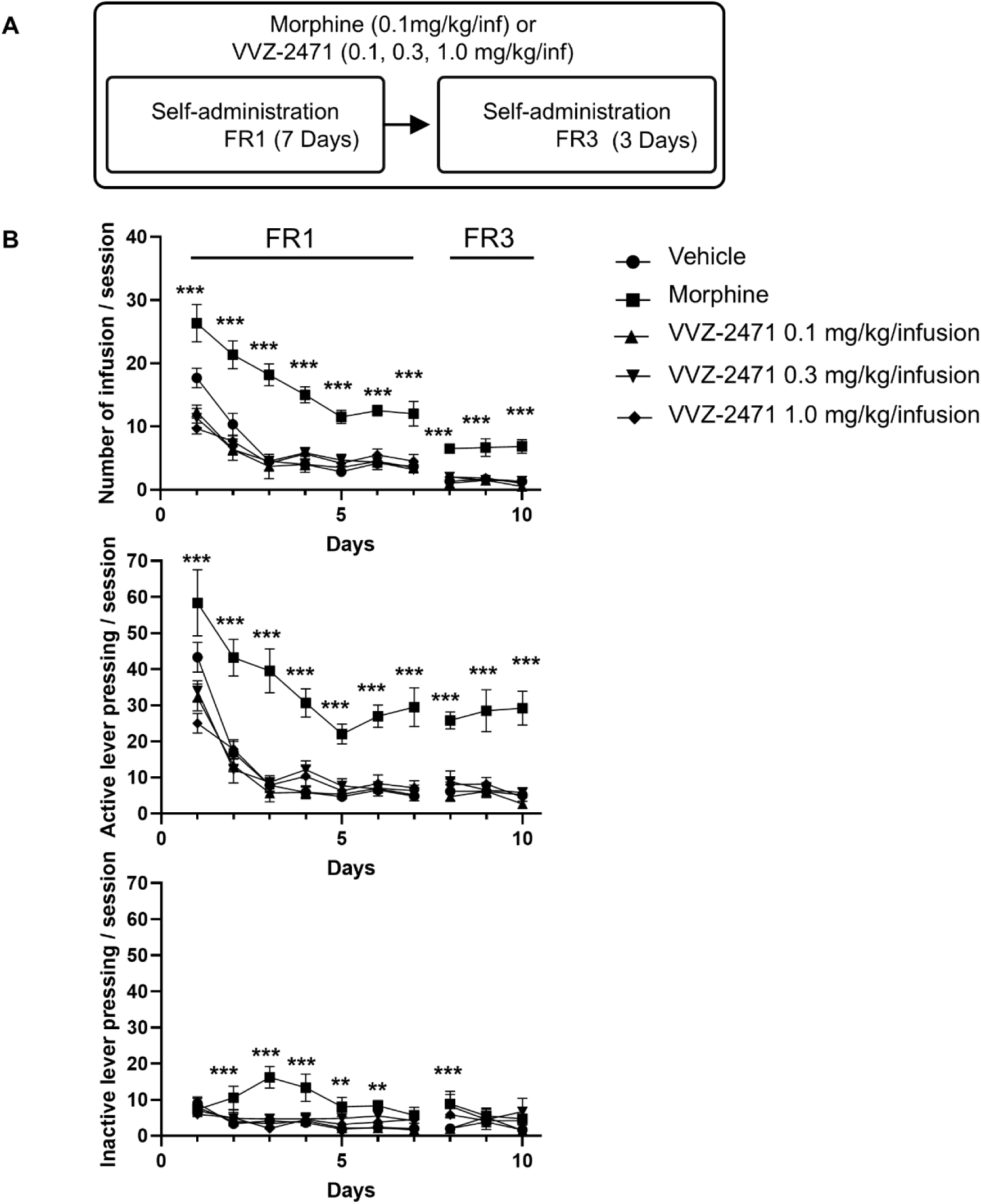
No abuse liability potential of VVZ-2471. **(A)** Schematic diagram of the experimental procedure for intravenous self-administration (IVSA) in rats. **(B)** the number of infusions (top), active lever presses (middle), and inactive lever presses (bottom) during 2 h sessions by vehicle (●), VVZ-2471 (0.1 (▴), 0.3 (▾), or 1.0 (◆) mg/kg/infusion), or morphine (▪) (0.1 mg/kg/infusion). (N=6 per each group). ** *P* < 0.01, *** *P* < 0.001 vs vehicle, Two-way ANOVA.

## 3. Discussion

Neuropathic pain remains a significant challenge in clinical pain management due to the complex mechanisms underlying peripheral and central sensitization. Recent efforts have focused on combination therapies targeting multiple pathways to enhance analgesic efficacy while minimizing adverse effects. However, combining multiple drugs to target these receptors presents significant challenges, as variations in the pharmacokinetic properties of individual drugs can result in short-lived effects and suboptimal therapeutic outcomes. Additionally, the concurrent use of multiple drugs increases the risk of drug-drug interactions, potentially leading to a higher incidence of adverse events ^49^. Previous research has demonstrated the efficacy of 5-HT2AR antagonism in pain relief, particularly when coupled with GlyT2 inhibition ^18–22^. GlyT2 inhibition elevates synaptic glycine concentrations, enhancing inhibitory neurotransmission and reducing neuronal excitability within pain pathways ^50^. In this current study, the GlyT2 component was replaced with a glutamatergic target, aiming to suppress excessive glutamatergic activity associated with neuropathic pain. Given the well-documented upregulation of mGluRs in chronic neuropathic and nociceptive pain ^12,39,51^, modulating mGluR5-related pathways presents a promising strategy for effective pain management. This led to the development of a novel multi-target therapeutic agent capable of antagonizing both 5-HT2AR and mGluR5 simultaneously-offering a potential solution to the limitations typically associated with combination therapies.

VVZ-2471, designed as an antagonist of 5-HT2AR (IC_50_ = 438.1 nM) and mGluR5 (IC_50_ = 87.1 nM) (**Fig. 2**), demonstrated robust analgesic effects comparable to those of gabapentin and morphine (**Fig. 4 and 5**). Notably, these effects were observed despite its moderate potency and at unbound free concentrations below its measured IC_50_ values (**Fig, 2 and Table 1**). Its pharmacokinetic (PK) profile supports sustained central nervous system (CNS) activity, exhibiting a high brain-to-plasma ratio of 1.87 at 2 hours post-administration and dose-dependent systemic exposure (**Table 1**). Furthermore, a strong PK/PD correlation (r² = 0.7379 in the SNL model, 0.5053 in the formalin model) reinforced its role in neuropathic pain and inflammation-induced pain modulation. Additionally, VVZ-2868, despite its lower potency for mGluR5, exhibited significant anti-allodynic and anti-nociceptive effects, reinforcing the therapeutic potential of dual-target modulation (**Fig. 4 and 5**).

Previous drug development efforts have focused on modulating glutamate signaling by targeting NMDA receptors. However, these single-target approaches have been substantially limited by adverse effects such as nausea, fatigue, and dizziness, which hindered their clinical applications ^30,31^. Given these limitations, we hypothesized that simultaneous antagonism of NMDA receptors and 5-HT2AR could enhance analgesic efficacy. Contrary to our expectations, the result revealed that the combination of MDL11,939 and radiprodil—a potent and selective antagonist of NR2B-containing NMDA receptors—failed to demonstrate sufficient efficacy in the SNL model (**Fig. S1**). This finding might emphasize the critical need for identifying optimal target combinations to maximize therapeutic benefits through target interactions.

Neurons in the superficial dorsal horn play a crucial role in pain information processing, particularly under pathological conditions characterized by increased mGluR5 and 5-HT2AR expression and heightened excitatory synaptic activity ^2,5,12,17^. Our findings further underscore the significance of 5-HT2AR and mGluR5 in pain signaling within this spinal region, supported by robust evidence demonstrating an increased frequency of sEPSC following the application of DHPG, 5-HT, and TCB-2, which was effectively suppressed by VVZ-2471 (**Fig. 3, S2 and S3**) ^16,17^. The anti-nociceptive effects of VVZ-2471 appear to be mediated via inhibition of ERK1/2 and AKT phosphorylation, following activation of 5-HT2AR and mGluR5, as well as in response to nociceptive stimuli like formalin (**Fig. 2 and 5**). These findings are consistent with recent studies demonstrating the role of 5-HT2AR and mGluR5 regulate pain signaling in the spinal dorsal horn through ERK1/2 and AKT phosphorylation ^52^.

While targeting the 5-HT2CR has shown potential benefits in treating schizophrenia and pain ^53,54^, it is also associated with serious adverse effects, including weight gain ^55–57^. VVZ-2471 demonstrated micromolar-range inhibitory potency for both the 5-HT2BR and the 5-HT2CR, suggesting a low likelihood of significant interaction with these receptors. Importantly, repeated administrations of VVZ-2471 over 14 days did not result in significant body weight changes, alleviating concerns about potential cross-activity with 5-HT2CR (**Fig. S4**). This finding suggests that any interaction between VVZ-2471 and the 5-HT2CR is insufficient to induce weight gain, further supporting its therapeutic safety profile.

A crucial aspect of VVZ-2471’s pharmacological profile is its high selectivity across molecular targets, minimizing off-target effects. Screening against 168 GPCRs, 371 kinases, and 87 ion channels and enzymes confirmed negligible interactions outside of its primary targets (**Fig. 2**). While minor binding affinity was detected for dopamine reuptake transporter (DAT) at high concentration (i.e., 10 μM VVZ-2471), the low unbound fraction (< 2% in humans) suggests a minimal risk of drug dependency (**Table 1**) ^58–60^. This hypothesis is further confirmed by direct intravenous infusion studies showing that VVZ-2471 does not induce drug dependence, even at doses of up to 1.0 mg/kg/infusion (**Fig. 7**).

The interplay between 5-HT2AR and mGluR5 presents a compelling therapeutic strategy for neuropathic pain management. Studies in mGluR5 knockout mice (mGluR5–/–) have revealed abnormal locomotor and exploratory behaviors, which were reversed by the 5-HT2AR antagonist M100907 ^61^. This finding suggests a compensatory regulation of serotonergic signaling in response to mGluR5 deficiency. Additionally, pharmacological inhibition of mGluR5 using MTEP (a potent mGluR5 antagonist) has been shown to increase 5-HT release, further highlighting the functional interplay between these receptor systems^62^. In inflammatory pain models, treatment with MPEP further amplified the increase in 5-HT levels following formalin or carrageenan injection ^63^, reinforcing the notion that mGluR5 antagonism may elevate serotonergic neurotransmission.

These compensatory dynamics imply that the simultaneous targeting of 5-HT2AR and mGluR5 may potentiate the analgesic effects of mGluR5 inhibition. Dual antagonism of 5-HT2AR and mGluR5 could mitigate neurotransmitter imbalance and lead to more robust and sustained pain relief. Our approach underscores the therapeutic potential of concurrently modulating these interconnected pathways to optimize treatment outcomes in neuropathic pain.

Numerous preclinical investigations have reported significant analgesic effects of 5-HT2AR and mGluR5 antagonists in female models. For example, selective 5-HT2A receptor blockade with sarpogrelate or ketanserin alleviated mechanical allodynia in lumbar disc herniation and diabetic neuropathy models in female rats ^64,65^. Similarly, mGluR5 inhibition with MPEP reduced post-surgical hypersensitivity in female rats ^66^. Together, these findings suggest that both receptor systems are critically involved in pain modulation irrespective of sex, reinforcing the translational potential of VVZ-2471 for male and female patients alike. Prospective comparative studies across sexes will further strengthen the clinical relevance.

Current first-line treatment options for neuropathic pain typically include tricyclic antidepressants (TCAs), serotonin-norepinephrine reuptake inhibitors (SNRIs), pregabalin, and gabapentin. However, their efficacy remains limited to approximately 40% of patients ^67^, leaving the majority reliant on opioid therapy. This dependence significantly increases the risk of opioid addiction, abuse, and other opioid-related complications. Our findings suggest that dual-target antagonism using VVZ-2471 may address critical gaps in neuropathic pain treatment, providing enhanced analgesic efficacy while reducing reliance on opioids (**Fig. 6**).

This could reinforce the case of dual 5-HT2AR and mGluR5 antagonists, such as VVZ-2471, as first-line treatment alternatives with minimal risk of tolerance development ^68,69^.

## 4. Materials and methods

### 4.1 Animals

Male Sprague–Dawley (SD) rats used for the spinal nerve ligation (SNL) and electrophysiology (5 weeks old) and formalin tests (8 weeks old) were purchased from Koatech Co., Ltd. (Pyeongtaek, Republic of Korea). Male SD rats used for intravenous self-administration experiments (7 to 8 weeks old) were purchased from Orient Bio Co., Ltd. (Seongnam, Republic of Korea). Prior to experimentation, all animals were maintained under controlled condition with a constant temperature (22 ± 3 ℃) and humidity (50 ± 10 %) in a 12-h light/dark cycle and free access to food and water. Handling and animal care were conducted in accordance with the guidelines of the Animal Care and Use Committee of the Institutional Animal Care and Use Committee of Vivozon Inc. (Gyeonggi-do, Republic of Korea) or Sungkyunkwan University (Gyeonggi-do, Republic of Korea). All animal experiments were approved by the institutional ethics committee (IACUC number: VVZ-IACUC-19R01004, 20R01004, 22R03012, 21R04002, 21R07006, 22R03011, 23R09007, SKKUIACUC2019-12-08-1).

### 4.2 Reagents

VVZ-2471 and VVZ-2868 were developed and synthesized by Research institute of Vivozon Inc. (Gyunggi-do, South Korea)^70^. 2-Methyl-6-(phenylethynyl) pyridine hydrochloride (MPEP), 5-hydroxytryptamine (5-HT), (4-bromo-3,6-dimethoxybenzocyclobuten-1-yl) methylamine hydrobromide (TCB-2), morphine, gabapentin, *α*-phenyl-1-(2-phenylethyl)-4-piperidinemethanol (MDL11,939), 7-(hydroxyimino)cyclopropa[*b*]chromen-1a-carboxylate ethyl ester (CPCCOEt), Fenobam and other reagents used for the *ex vivo* and *in vivo* experiments were purchased from Tocris Bioscience (Bristol, UK) and Sigma–Aldrich (MO, USA) unless otherwise specified. Morphine sulfate hydrate for *in vivo* experiments was purchased from the Ministry of Food and Drug Safety (MFDS) of the Republic of Korea. Rabbit anti-phospho-p44/42 MAPK (ERK1/2, Thr202/Tyr204), rabbit anti-p44/42 MAPK (ERK1/2), rabbit anti-phospho-AKT (Ser473), rabbit anti-AKT and anti-rabbit IgG were purchased from Cell Signaling Technology (MA, USA). Other reagents and materials for *in vitro* experiments were purchased from ThermoFisher Scientific (MA, USA), Corning Inc. (NY, USA), and Bio-RAD Laboratories (CA, USA). Verapamil was purchased from Sigma–Aldrich (MO, USA).

### 4.3 5-HT2AR overexpression cell (HEK293T cell) and IP1 assay

HEK293 cell line, stably transfected with the pCi/neo vector (Promega Biotechnology Company, WI, USA) containing the coding sequence for the human 5-HT2A receptor (BC074849, P28233), was obtained from Cerep (France). The cells were maintained in 88.4% Dulbecco’s Modified Eagle Medium (DMEM) supplemented with GlutaMAX and sodium pyruvate, 10% of dialyzed fetal bovine serum (FBS) (ThermoFisher Scientific, MA, USA), and 1.6% or 800 μg/mL Geneticin. 15 nM of 5-HT (approximately EC_80_) was applied to the cells preincubated with a series of concentrations of VVZ-2471 for 10 minutes at room temperature, followed by incubation for 30 minutes at 37 °C in a dark place. IP 1 generation *in vitro* was calculated using fluorescence intensity ratio changes at 665 nm and 620 nm detected with SpectraMax i3x (Molecular Devices, LLC, CA, USA) according to the Homogeneous Time Resolved Fluorescence (HTRF) assay method. IC_50_ value was determined by nonlinear regression analysis using GraphPad Prism9.

### 4.4 Target selectivity

G protein-coupled receptor (GPCR) panel profiling for 168 targets was performed by DiscoverX (gpcrMAX℠) (DiscoverX, CA, USA) to evaluate the potential interaction of VVZ-2471 with various GPCRs. A kinase assay panel comprising 371 different kinases was conducted in the Reaction Biology (PA, USA). The SafetyScreen 87 panel assay was performed at EurofinsCerep (France) using validated radioligand competition binding assays and enzymatic inhibition assays. Detailed experimental conditions for each of these 87 assays are available on the Eurofins website (www.eurofinspanlabs.com). Subtype selectivity experiments for mGluR7 and mGluR8 were conducted using the metabotropic glutamate receptors panel service (EuroscreenFast, Belgium). mGluR1 functional assay was conducted with GPCRscan service (DiscoverX, CA, USA) *via* PathHunter *β*-arrestin assays. Additionally, subtype selectivity experiments for the serotonin 2B receptor (5-HT2BR) and serotonin 2C receptor (5-HT2CR) were conducted using Gq cellular functional assays (EurofinsCerep, France).

### 4.5 Primary neuronal culture

Rat brains were dissected out from embryonic Day 17, and the cerebral cortices were quickly isolated at 4 °C in Hanks’ Balanced Salt Solution (HBSS) supplemented with 4-(2-hydroxyethyl)-1-piperazineethanesulfonic acid (HEPES) buffer. These tissues were treated with HBSS containing Accutase solution and 25 μg/mL of DNase Ι at 37 °C for 30 minutes, resulting in a single-cell suspension in culture medium supplemented with B27 and 1% penicillin–streptomycin in Neurobasal medium. The dissociated cells were plated in a poly-D-lysine-coated 6-well plate (Corning Inc., NY, USA) at a density of 1.2 × 10^6^ cells per well and cultured in humidified air with 5% CO_2_ at 37 °C for 14 days prior to drug treatment. MPEP, MDL11,939, Fenobam, and VVZ-2471 were preincubated for 20 minutes before the addition of DHPG or TCB-2. CPCCOEt was included in the medium 20 minutes before drug treatment to inhibit mGluR1 activation.

### 4.6 Western blot

Extracts were obtained from primary cortical cultures and rat spinal cord dorsal horn using RIPA lysis buffer containing 0.5 M of ethylene-diamine-tetraacetic acid (EDTA), along with halt protease and phosphatase inhibitor cocktail (ThermoFisher, MA, USA). For spinal cord tissue, the dorsal horn of the lumbar spinal cord was harvested 20 minutes after formalin injection, followed by homogenization in ice-cold T-PER^TM^ lysis buffer. The homogenates were then centrifuged at 14,000×*g* at 4 °C for 15 minutes, and the protein concentration of the supernatant was measured using a Bradford assay. Equal amounts of proteins were resolved in 4 to 12% Bis-Tris gel electrophoresis (Bio-RAD Laboratories, CA, USA) and subsequently transferred to a polyvinylidene difluoride (PVDF) membrane (Bio-RAD Laboratories, CA, USA). Following membrane blocking in Tris-buffered saline (TBS) containing 5% skim milk, antibody incubation was conducted. The primary antibodies used were rabbit anti-phospho-p44/42 MAPK (ERK1/2, Thr202/Tyr204, 1:1,000), rabbit anti-p44/42 MAPK (ERK1/2, 1:1,000), rabbit anti-phospho-AKT (Ser473, 1:1,000), and rabbit anti-AKT (1:1,000). Anti-rabbit HRP IgG was used as the secondary antibody (1:2,000). Detection and quantification were performed by ChemiDoc system (Bio-RAD Laboratories, CA, USA) with enhanced chemiluminescence.

### 4.7 Slice preparation

The preparation of spinal cord slices was carried out as previously described ^71^. Before anesthetization, *N-*methyl-D-glucamine (NMDG) solutions (aerated with CO_2_/H_2_O for 30 minutes) were pre-chilled in a slush condition, and artificial cerebrospinal fluid (ACSF) was also aerated with CO_2_/H_2_O. The vibratome chamber was pre-chilled at –20 ℃ until an icy layer became visible. Animals were deeply anesthetized *via* intraperitoneal injections of urethane (1.5 g/kg). A lumbosacral laminectomy was performed, and a segment of the spinal cord (approximately 2.0 cm) along with the dorsal roots was placed in an ice-cold NMDG solution. This spinal cord segment was then affixed to an agar block and mounted on a cutting stage, ensuring that the longitudinal axis of the spinal cord was perpendicular to the razor blade of a VT1200 Vibratome (Leica Biosystems, IL, USA). Slices with a thickness of 300 to 400 μm were prepared. After a recovery period of 30 minutes in a warmed chamber, the spinal cord slices were maintained at room temperature in aerated ACSF until further use.

### 4.8 Electrophysiology

The composition of the solutions is detailed below: NMDG cutting solution (135 mM NMDG, 1 mM KCl, 1.2 mM KH_2_PO_4_, 1.5 mM MgCl_2_, 0.5 mM CaCl_2_, 20 mM choline bicarbonate, and 10 mM glucose), ACSF (130 mM NaCl, 24 mM NaHCO_3_, 3.5 mM KCl, 1.25 mM NaH_2_PO_4_, 1 mM MgCl_2_, 2 mM CaCl_2_, and 10 mM glucose, aerated with 95% O_2_/5% CO_2_), and internal solution (108 mM Cs-Methane sulfonate, 4 mM MgCl_2_, 1 mM EGTA, 9 mM HEPES, 5 mM Mg-ATP, 15 mM phosphocreatine (di-) TRIS, 1 mM Na-GTP, and 5 mM QX-314, pH 7.4, 280 mOsm).

Whole-cell patch-clamp recordings were conducted on neurons in the superficial dorsal horn of the spinal cord. The recording pipette (2 to 4 MΩ) contained a cesium (Cs+)-based internal solution. Spontaneous excitatory postsynaptic currents (sEPSCs) were recorded at a holding potential of –70 mV, where the equilibrium potential for chloride is close to 0 mV. The absence of inhibitory post-synaptic currents (IPSCs) was confirmed by periodically applying cyanquixaline (CNQX, 10 μM, Tocris, USA) and D-AP5 (50 μM) (Tocris, USA). All recordings were conducted at a temperature range of 29.5 to 30 ℃ using a CL-200A heater-cooler controller (Warner Instrument, CT, USA) in a chamber with a constant flow of bath solutions. Synaptic responses were recorded using a MultiClamp 700B amplifier (Axon Instruments, CA, USA), digitized with a Digidata 1550B (Axon Instruments, CA, USA), and data acquisition was performed by Clampex 11.0.3 (Axon Instruments, CA, USA). Data analysis was conducted using Clampfit software. DHPG, 5-HT, and TCB-2 were bath-applied with a constant flow to induce an increase in sEPSC after establishing a stable basal level of synaptic responses. VVZ-2471, MPEP, MDL11,939, and Basimglurant were applied 5 minutes after each agonist application. The amplitude and frequency of sEPSC were analyzed over a consecutive 30-seconds duration.

### 4.9 In vivo studies for SNL and formalin-induced pain models

All tests and analysis were conducted in a blinded manner with respect to the experimental conditions. Surgical procedures were performed in accordance with a previous study for the SNL model ^25^. Animals received the SNL surgery at the terminals of the dorsal root ganglion on the left side of the lumber 5 and 6, and the nerve was securely tied with a suture thread at the entrance to the sciatic nerve. Following the closure of the incision, the rats were allowed to recover for 2 weeks.

Pain measurements were assessed using a von Frey filament of 0.41 to a maximum of 15.8 g, applied to the plantar surface of the ipsilateral hind paws. The paw withdrawal threshold (PWT), characterized by nocifensive behaviors—including rapid paw withdrawal, licking, or shaking of the paw—either during stimulus application or immediately after filament removal, was recorded according to the up- and-down method as previously described ^72,73^. Each measured PWT value, a logarithmic gram value of measured unit of force, was converted to a percent reversal (% reversal) using the following equation:

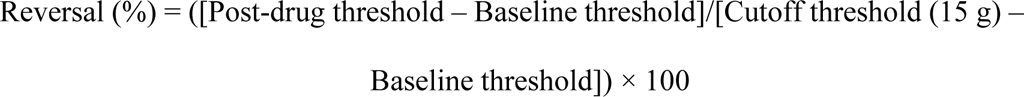

where post-drug threshold means the PWT after drug injection.

Randomization was conducted based on the results of the von Frey test to ensure that each test group exhibited no significant differences in their baseline PWT values. MDL11,939, MPEP, and a combination of both drugs were subcutaneously administered 0.5 hours prior to pain measurement. Gabapentin (Combi-Blocks, CA, USA) was administered intraperitoneally 1 hour before, while VVZ-2471 and VVZ-2868 were given orally 1 to 2 hours prior to pain measurement.

For the formalin test, animals were randomized based on their body weights and transferred to the testing room at least 1 h prior to the formalin injection. After a habituation period, a 3% formalin solution (50 μL) was injected into the plantar surface of the left hind paw. Pain behavior was assessed by recording the frequency of flinching and the duration of licking from 15 to 40 minutes at 5-minute intervals ^26^. MDL11,939, MPEP, and a combination of both drugs were subcutaneously administered 30 minutes before the formalin injection. Morphine was administered subcutaneously 10 minutes prior, while VVZ-2471 and VVZ-2868 were given orally 100 minutes before the formalin injection.

MDL11,939 and MPEP were dissolved in a mixture of *N,N*-dimethylacetamide (DMA) (Sigma–Aldrich, MO, USA) and propylene glycol (PG) in a 2:8 ratio. Gabapentin was dissolved in phosphate buffered saline (PBS), and morphine sulfate was dissolved in 0.9% saline.

### 4.10 Intravenous self-administration (IVSA) study

IVSA tests were conducted with minor modifications to the previously reported experimental design^74^. Animals were trained in operant chambers (28 cm × 26 cm × 20 cm) (Med Associates, Inc., St. Albans, VT, USA) equipped with response levers and cue lights. Drugs were delivered *via* an intravenous catheter connected to a syringe pump (Razel Scientific Instruments, Georgia, VT, USA), which was positioned on top of the cubicle. All experimental sessions were controlled and recorded by a PC using a custom interface and software within the experimental room. Following randomization, IVSA of the drugs was conducted under a fixed ratio (FR)1 schedule lasting 2 hours for 7 consecutive days. Each response on the active lever resulted in a drug infusion (0.1 mL over 4 seconds) followed by a 20-second timeout with light cues. The schedule was subsequently increased to FR3 for an additional 3 days. Inactive lever presses were recorded but had no programmed consequences.

### 4.11 Pharmacokinetic analysis

Plasma samples were prepared from 0.5 mL of blood collected in heparin-coated 1.5 mL-Eppendorf tubes, which were centrifuged for 5 minutes at 5,000×*g* at room temperature. Rat brain tissues were homogenized in a two-fold volume (*w*/*v*) of PBS using a FastPrep-24^TM^ 5G (MP Biomedicals, USA) equipped with ceramic beads. From this homogenate and plasma samples, 50 µL was combined with acetonitrile containing verapamil (an internal standard) to facilitate protein precipitation. Following centrifugation at 5,900×g for 10 minutes at 4 °C, the supernatant was analyzed using the LC-MS/MS method.

Chromatographic analyses were conducted using a Unison UK-18 (Imtakt, Japan) C18 column (75 mm × 2.0 mm, 3 µm) at a temperature of 40 °C. The chromatography eluates were analyzed with the SCIEX QTRAP 4500 LC-MS/MS system (AB SCIEX, MA, USA). Data collection was performed using Multiquant software (3.0.2). Liver microsomal stability and plasma protein binding were assessed as previously described ^75,76^. The terminal half-life (*t*_1/2_), volume of distribution at elimination phase (*V*_z_), clearance (CL), and area under the plasma concentration–time curve (AUC) were calculated using noncompartmental analysis with Phoenix® WinNonlin® software (8.3.5.340) (Certara, St. Louis, MO, USA). The values for peak time (*T*_max_) and peak concentration (*C*_max_) were obtained directly from the original data.

### 4.12 Statistical analysis

All data were expressed as the mean ± standard error of the mean (S.E.M.) with exception of pharmacokinetic experiments, which were expressed as the mean ± standard deviation (S.D.). For pain assessment in the SNL and formalin models, all data were analyzed using GraphPad Prism 9. A one way or two-way ANOVA, followed by Tukey’s multiple comparisons test, was employed for analysis in the SNL and formalin tests. For the electrophysiological data, spontaneous events were automatically detected using the event detection function in Clampfit (11.0.3), and were subsequently examined manually in a post-hoc analysis to exclude false-positive events. A Student’s t-test was used for the statistical analysis of synaptic responses and western blot results. For the self-administration test, statistical analyses were conducted using a two-way ANOVA followed by Fisher’s LSD *post hoc*. *P* < 0.05 was considered statistically significant. In this study, ‘n’ represents the number of data points or *ex vivo* slices, while ‘N’ represents the number of animals or independent culture sets derived from different litters. No power analyses were used to determine sample size, but our sample sizes are similar to those from previous studies ^77–80^.

## Author contributions

Daekyu Choi, Hyun Jin Heo, Hanmi Lee conceptualized the study. Daekyu Choi supervised drug chemistry. Hyun Jin Heo conducted and analyzed *in vivo* experiments. Jayzoon Im and Hanmi Lee conducted and analyzed *ex vivo* electrophysiology experiments. Haeyoung Shin and Ah Hyun Kim conducted and analyzed *in vitro* experiments. Geonho Lee conducted and analyzed pharmacokinetics experiments. Kwang-Hyun Hur and Choon-Gon Jang conducted IVSA experiments. Yoonmi Nho participated in conceptualization of the draft with substantial intellectual contribution. Hanmi Lee supervised entire study. Hanmi Lee mainly wrote, and Hanmi Lee and Yoonmi Nho edited the manuscript. All authors participated in the discussions and in writing the manuscript, and approved the final version.

## Acknowledgements

We thank Drs. Jin Mo Chung and Jun-Ho La, Department of Neurobiology, University of Texas Medical Branch, Galveston, Texas for reading the manuscript and making valuable suggestions. This research was supported by the Basic Science Research Program (2022R1A6A1A03054419) through the National Research Foundation of Korea to CGJ.

## Availability of Data and Reagents

The data and reagents used in this study are available from the corresponding author upon reasonable request.

## Supplemental figure legends

**Figure S1.**
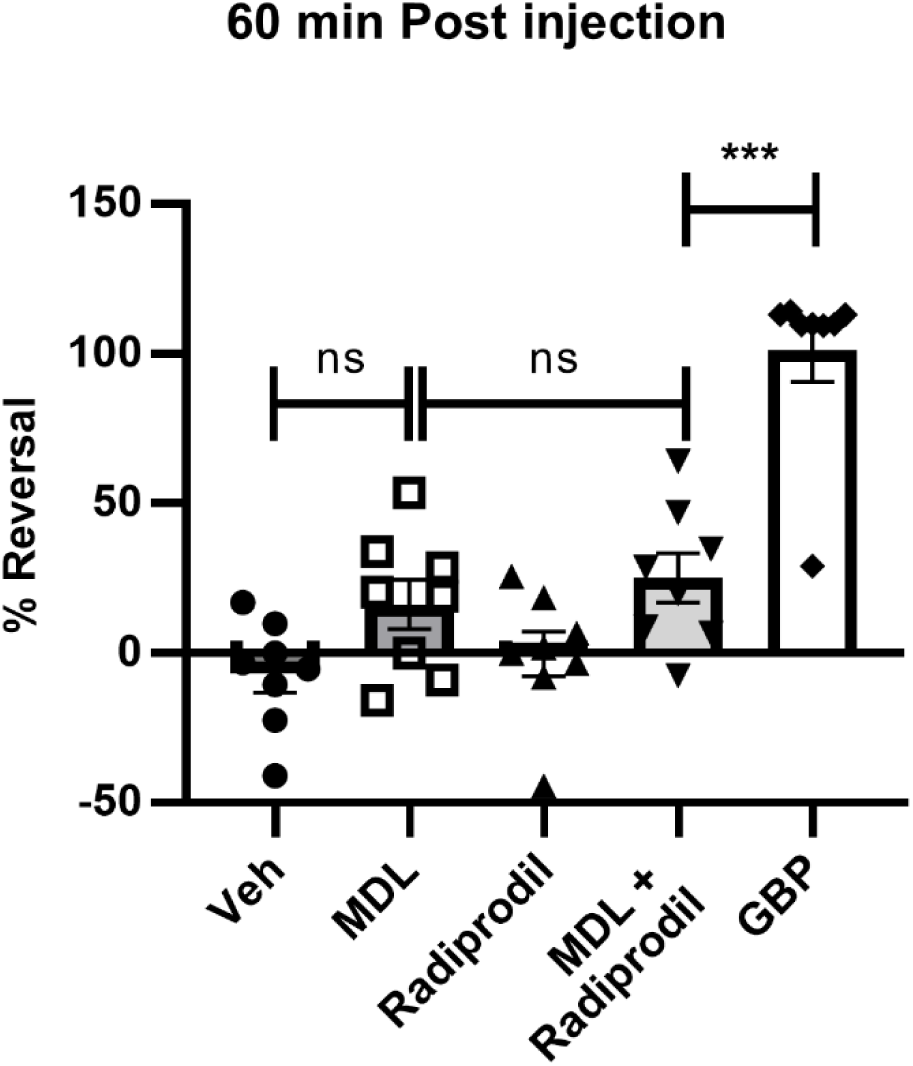
No efficacy enhancement of dual antagonism against 5-HT2AR and NMDA receptor in SNL model. Percent (%) reversal of the paw withdrawal threshold in the SNL model after treatment of vehicle (2:8 DMA/PG, s.c.), MDL (5 mg/kg, s.c.), Radiprodil (30 mg/kg, s.c.), a combination of MDL and Radiprodil, or GBP (65 mg/kg, i.p.) (N = 8 rats per group). Error bars present S.E.M., “ns” means no significance (*P* > 0.05); \*\*\**P* < 0.001, One-way ANOVA.

**Figure S2.**
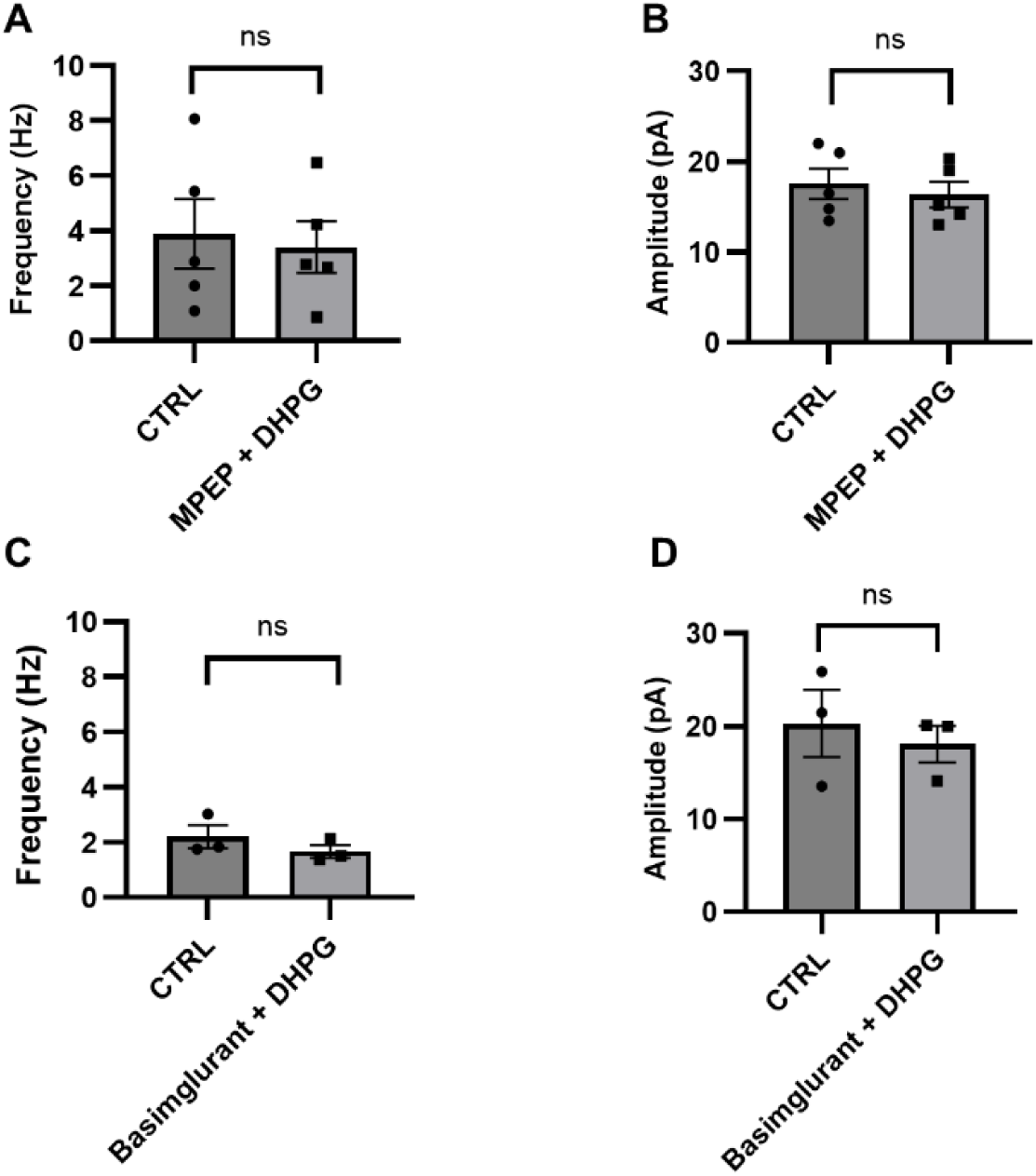
DHPG-induced sEPSC in the presence of MPEP and Basimglurant. **(A)** Frequency (Hz) changes in the presence of MPEP (10 μM) and MPEP plus DHPG (3 μM), Control (CTRL): 3.9 ± 1.1; MPEP+DHPG: 3.4 ± 0.8, n=5 / N=1). **(B)** Amplitude (pA) changes in the presence of MPEP and MPEP plus DHPG (3 μM) (CTRL: 17.6 ± 1.5; DHPG + MPEP: 16.3 ± 1.3, n=5 / N=1). **(C)** Frequency changes in the presence of Basimglurant (10 μM) and Basimglurant plus DHPG (3 μM) (CTRL: 2.2 ± 0.3; DHPG + Basimglurant: 1.7 ± 0.2, n=3 / N=1). **(D)** Amplitude changes in the presence of Basimglurant (10 μM) and Basimglurant plus DHPG (3 μM) (CTRL: 20.3 ± 2.9; DHPG + Basimglurant: 18.1 ± 1.6, n=3 / N=1). Error bars present S.E.M., “ns” means no significance (*P* > 0.05); \**P* < 0.05; \*\**P* < 0.01; \*\*\**P* < 0.001, Two-tailed t-test.

**Figure S3.**
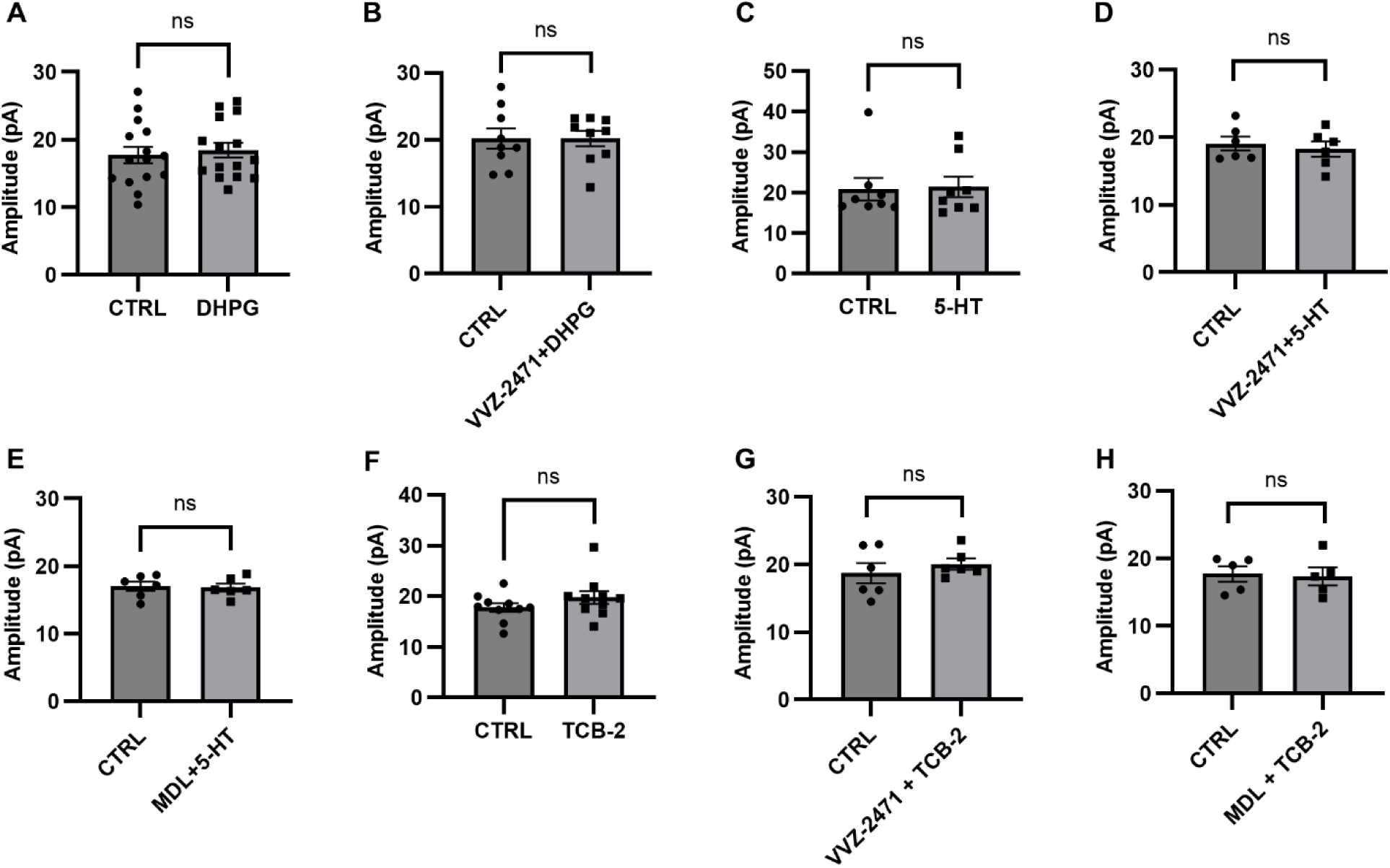
No significant effects of VVZ-2471 on the amplitudes of sEPSC. **(A)** Amplitude (pA) changes by DHPG alone (3 μM, Control (CTRL): 17.7 ± 1.2; DHPG: 18.5 ± 1.1, n=15 / N=9). **(B)** Amplitude changes by DHPG in the presence of VVZ-2471 (10 μM) (CTRL: 20.2 ± 1.4; DHPG + VVZ-2471: 20.1 ± 1.1, n=9 / N=3). **(C)** Amplitude changes by 5-HT (50 μM, CTRL: 20.8 ± 2.6; 5-HT: 21.4 ± 2.3, n=8 / N=6). **(D)** Amplitude changes by 5-HT in the presence of VVZ-2471 (CTRL: 19.1 ± 1.0; 5-HT + VVZ-2471: 18.2 ± 1.0, n=6 / N=3). **(E)** Amplitude changes by 5-HT in the presence of MDL11,939 (MDL) (10 μM, CTRL: 17.0 ± 0.6; 5-HT + MDL: 16.8 ± 0.5, n=6 / N=4). **(F)** Amplitude changes by TCB-2 alone (100 μM, CTRL: 17.7 ± 0.8; TCB-2: 19.7 ± 1.2, n=10 / N=5). **(G)** Amplitude changes induced by TCB-2 in the presence of VVZ-2471 (10 μM, CTRL: 18.7 ± 1.3; TCB-2 + VVZ-2471: 20.1 ± 0.7, n=6 / N=3). **(H)** Amplitude changes induced by TCB-2 in the presence of MDL (10 μM, CTRL: 17.5 ± 1.2; TCB-2 + MDL11,939: 17.4 ± 1.3, n=5 / N=3). Error bars present S.E.M., “ns” means no significance (*P* >0.05), Two-tailed t-test.

**Figure S4.**
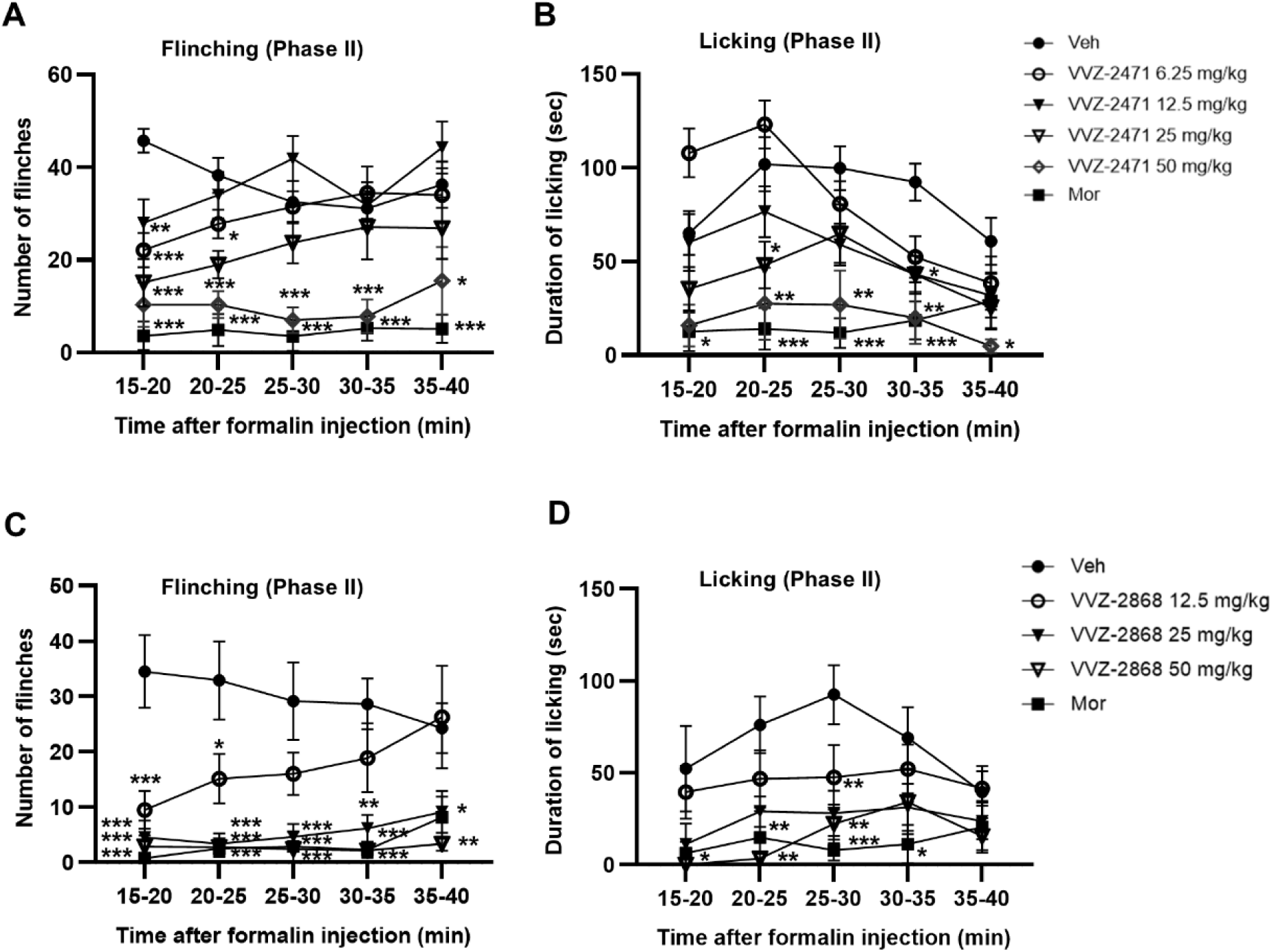
Anti-nociceptive effects of dual antagonists against 5-HT2AR and mGluR5 in the formalin-induced pain model. Time-course of flinches **(A, C)** and licking **(B, D)** responses induced by formalin injection to the rats treated with VVZ-2471 (p.o.), VVZ-2868 (p.o.), or Morphine (MOR, 2 mg/kg, s.c.). N = 8 rats per group. Error bars present S.E.M., \**P* < 0.05; \*\**P* < 0.01; \*\*\**P* < 0.001 *vs* Vehicle, Two-way ANOVA.

**Figure S5.**
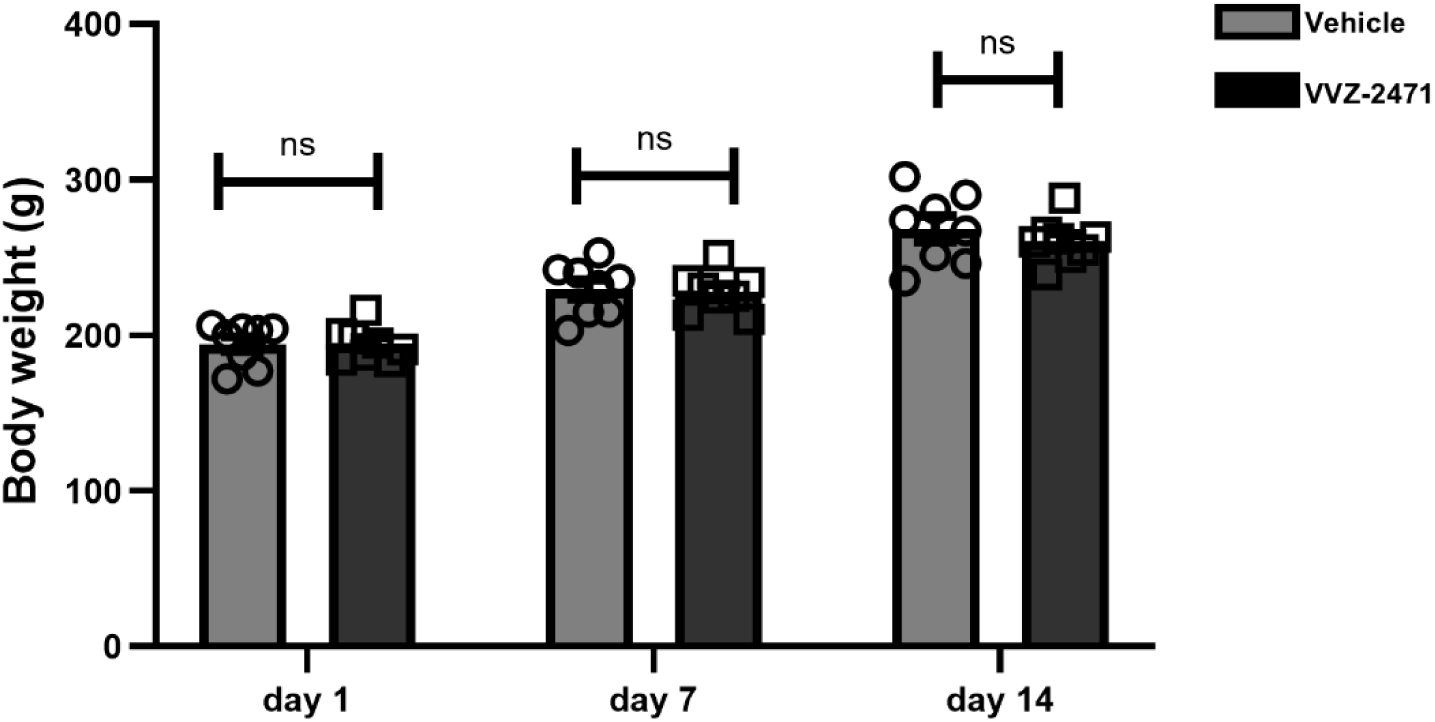
Effect of VVZ-2471 on body weight. VVZ-2471 (25 mg/kg, p.o.) and vehicle (p.o.) were administered once a day for 14 consecutive days (N=8 rats per group). Body weight measured on Days 1, 7, and 14. Error bars present S.E.M., “ns” means no significance (*P* > 0.05), One-way ANOVA.

